# A data-driven modeling framework for mapping genotypes to synthetic microbial community functions

**DOI:** 10.1101/2025.01.04.631316

**Authors:** Yili Qian, Sarvesh D. Menon, Nick Quinn-Bohmann, Sean M. Gibbons, Ophelia S. Venturelli

## Abstract

Microbial communities play a central role in transforming environments across Earth, driving both physical and chemical changes. By harnessing these capabilities, synthetic microbial communities, assembled from the bottom up, offer valuable insights into the mechanisms that govern community functions. These communities can also be tailored to produce desired outcomes, such as the synthesis of health-related metabolites or nitrogen fixation to improve plant productivity. Widely used computational models predict synthetic community functions using species abundances as inputs, making it impossible to predict the effects of species not included in the training data. We bridge this gap using a data-driven community genotype function (dCGF) model. By lifting the representation of each species to a high-dimensional genetic feature space, dCGF learns a mapping from community genetic feature matrices to community functions. We demonstrate that dCGF can accurately predict communities in a fixed environmental context that are composed in part or entirely from new species with known genetic features. In addition, dCGF facilitates the identification of species roles for a community function and hypotheses about how specific genetic features influence community functions. In sum, dCGF provides a new data-driven avenue for modeling synthetic microbial communities using genetic information, which could empower model-driven design of microbial communities.

## INTRODUCTION

Microbial communities are important drivers of ecosystems on Earth spanning the human gut microbiome to the soils [1]. They perform a diverse repertoire of biochemical and physical transformations (i.e., functions), such as degradation of dietary substrates or production of metabolites, that alter the environment they reside in. Leveraging these capabilities, there has been an increasing interest to assemble defined, synthetic microbial communities from bottom-up to perform target functions that address important societal challenges in health [2], environment [3], [4], agriculture [3], [5], and sustainable bioprocessing [6] (**Figure 1**A). For example, microbial communities have been designed to replace fecal microbiota transplantation for treating recurrent *C. difficile* infection [7], modulate human immune response [8], inhibit fungal plant pathogens [9], and enhance plant nitrogen fixation for increased agricultural yield [10]. Unlike microbiome samples collected from the environment, where community composition is unknown *a priori* and environmental conditions vary across samples, synthetic microbial communities are assembled in defined lab environments with designer-specified initial composition. This allows researchers to quantitatively study the impact of particular species or environmental factors on community assembly and functions, which may be extendable to natural microbiomes [11]–[23].

**Figure 1:**
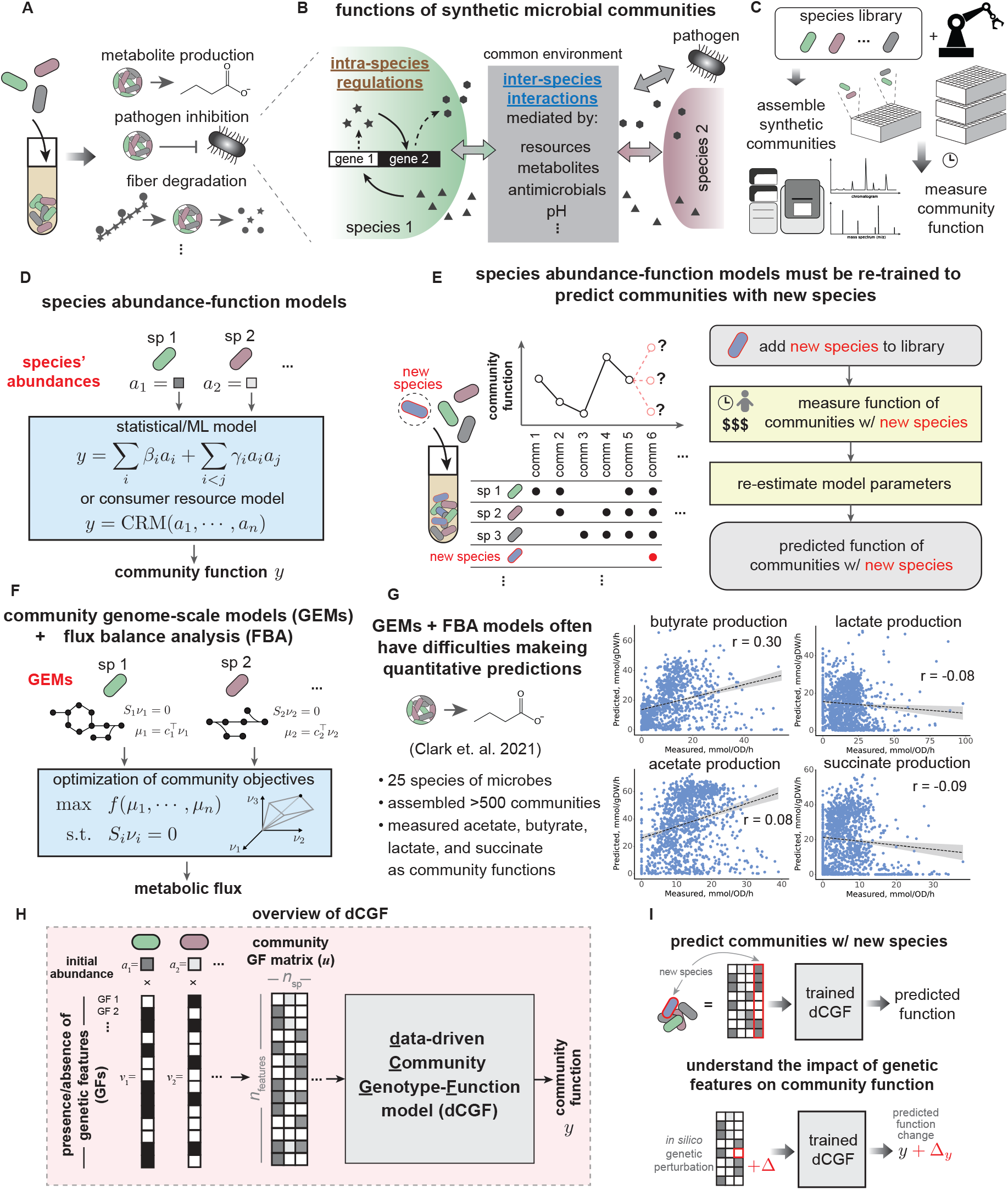
Prediction of synthetic microbial community functions. (A) Example functions of synthetic microbial communities include the rates at which they can produce metabolites of interests, degrade complex fibers into simple sugars, and their ability to inhibit pathogens. (B) Functions of microbial communities are affected by intra-species regulations and inter-species interactions. The ability of a synthetic community to inhibit a pathogen, for instance, is determined by the common environment the community members shape. The common environment is characterized by, for example, pH, the concentrations of different types of nutrients that support pathogen growth, and the anti-microbials produced by community members to inhibit the pathogen. Each community member senses the common environment to regulate gene expression, creating intra-species regulations that regulate the production or consumption of molecular effectors, including resources, metabolites, or antimicrobials. Variations in the concentrations of these molecular effectors in the common environment affects the growth and metabolic states of other species in the community, giving rise to inter-species interactions. (C) Lab automation has been applied to assemble hundreds of synthetic communities from a library of species in one high-throughput experiment. The communities functions are measured after the assembled communities grow for a fixed amount of time. (D) Species abundance-function models (SAMs) are one of the most common types of mathematical models used to quantify and subsequently predict community functions. SAMs map the abundance (or presence/absence) of species in a synthetic community to the community’s function. Both data-driven (i.e., statistical or machine learning models) and mechanistic models (e.g., consumer resource models) have been used. (E) A trained SAM cannot be used to predict the function of a community that includes a new species absent in the training data. In order to predict the function of such a community, one has to measure the functions of some communities including the new species as new training data, and then re-estimate model parameters. This is often a costly process. (F) Applying flux balance analysis (FBA) on community genomes-scale models (GEMs) is a common approach to predict metabolic functions of synthetic microbial communities. This approach does not rely on any training data. Instead, it takes advantage of each species’ known genome annotations and assumes that the assembled community optimizes an objective function, often associated with maximizing biomass. (G) GEMs+FBA was used to predict the rates at which synthetic human gut microbial communities produce health-related metabolites, including acetate, butyrate, lactate, and succinate (see **Methods**). Scatter plots show correlation between predicted and measured metabolite production rates. Measured data were obtained from experimental study in [27], where over 500 synthetic microbial communities were assembled from a set of 25 species. (H) Data-driven community genotype-function model (dCGF) takes the community genetic feature (GF) matrix *u* as inputs to predict community functions *y* as outputs. The community GF matrix is composed of the binary vectors of each species’ genetic features (*v*_*i*_) scaled by its initial relative abundance (*a*_*i*_). (I) A trained dCGF can predict the function of communities including new species absent in training data, as long as the GFs of the new species are known and each GF is present in at least one species in traing data. By perturbing elements in the community GF matrix, a trained dCGF can also be used investigate the impact of GFs on community function.

The combinatorial nature of microbial communities is a major bottleneck to its bottom-up design and analysis. For example, microfluidic systems can assemble up to 10^7^ defined communities per experiment [24], [25]. Yet, this number still pales in comparison to more than 10^15^ possible communities that can be assembled from, for example, 50 candidate species. To overcome this formidable barrier, computational models are becoming essential to capture complex biological processes and to efficiently navigate the community-function landscape, which is the mathematical relationship mapping community composition to function [26]. For example, models have been developed to predict health-relevant metabolite profiles from initial species abundances. These models have been used to identify specific communities with enhanced production of the health-beneficial metabolite butyrate [27]–[29].

The function of a synthetic microbial community is determined by complex intra-cellular regulatory networks and inter-species interactions (**Figure 1**B) that are often not well-characterized and understood. Consequently, mechanistic mathematical models often have limited success in small communities composed of well-characterized species and environments [23], [29], [30]. Statistical and/or machine learning (ML) models have the flexibility to learn unknown regulations, interactions, and community assembly rules from training data [28], [31]–[33]. These data-driven models leverage high-throughput lab automation techniques [18], [34], which allow researchers to quantify the functions of hundreds to thousands of defined synthetic communities assembled from a set of species used for model training (**Figure 1**C), which we call training species. However, existing data-driven models use initial abundances or presence/absence (i.e., one-hot encoding) of training species as inputs to predict community functions. Consequently, a model trained on a given dataset cannot be reused to predict communities containing different species that are absent in training data in the same environmental context. Developing this capability could enable the selection of specific new species beyond the set of training species to optimize a target function using experimental design techniques [31].

In contrast to natural microbiome data, where species composition needs to be inferred from metagenomic or 16S rRNA sequencing data, synthetic microbial communities can be assembled from the bottom-up in a defined environment using species with known genetic information (e.g., genome sequence) and initial abundances. This property makes it possible to infer how adding different species can impact a synthetic community’s function. For instance, the genome of the newly added species may share many genetic features (GFs) already present in species in training data. This makes it possible to predict the impact of the newly added species on a community metabolic function based on its “metabolically similarity” to those already present in training data. Recent studies have shown that the metabolic functions of single species [35] as well as the metabolic function of microbial communities can both be predicted from genetic information [36]. For example, a mechanistic consumer-resource model (CRM) was used by Gowda et al [36] to represent the dynamics of denitrification in synthetic soil microbial communities. This study used a linear regression model to map the presence/absence of genes to each species’ CRM parameters, such as resource consumption rates. Therefore, a model that uses genetic information as inputs can uncover how these genetic features quantitatively contributes to community functions.

Here we propose a data-driven community genotype-function (dCGF) modeling framework to map the genetic features (GFs) of species to a community function in a fixed environmental context using tailored ML models. dCGF has a more flexible structure in comparison to previous models mapping genetic information to community properties [36]. Specifically, compared to the model in [36], dCGF can account for inter-species interactions beyond CRM and use nonlinear relationships to quantify the impact of GFs’ presence/absence on community function. The species genotype to community function mapping learned by dCGF from training data can be re-used to predict the functions of new communities assembled partly or entirely from species beyond the initial set of training species *in silico*, thus eliminating the need to characterize these communities experimentally. We evaluate the prediction performance of candidate realizations of dCGF using synthetic data generated from mechanistic model simulations, which include communities with multiple modes of complex inter-species interactions and intra-cellular regulations. Using published experimental data, we demonstrate that dCGF can not only predict functions of communities assembled from the training species (i.e., interpolation), but also communities assembled partially or completely from species absent in the training data (i.e., extrapolation) in the same environmental context. Using sensitivity analysis, a trained dCGF can guide hypotheses about GFs that may contribute to a given function. Finally, we demonstrate that the model provides a data-driven approach to reveal species’ contributions to a particular community function. dCGF is a proof-of-concept data-driven modeling framework that learns species genotype to community function mappings in synthetic microbial communities. By re-using learned genotype-function mapping to species absent from training data, This modeling framework can substantially improve model re-usability to benefit model-guided design and understanding of the complex behaviors of microbial communities.

## RESULTS

### Existing modeling frameworks have limitations in predicting synthetic microbial community functions

Current computational models to predict synthetic microbial community functions include species abundance-function models (SAMs) and community genome-scale models (GEMs). SAMs predict community function from species’ initial abundances (**Figure 1**D). SAMs can take the form of data-driven models, including statistical regression and machine learning models [28], [31]–[33], or mechanistic models, including consumer resource models [12], [14], [22] and dynamic ecological models, such as the generalized Lotka-Volterra (gLV) model [18], [21]. Although SAMs have demonstrated the ability to accurately predict functions of communities assembled from a set of training species, they cannot predict communities that include new species absent from the set of training species (**Figure 1**E). Therefore, when introducing a new species into the synthetic community, a SAM must be re-trained on new experimental data of communities containing this species, thus requiring additional time and cost.

Alternatively, using constraint-based modeling (e.g., flux balance analysis, FBA), the metabolic fluxes of a microbial community can be predicted using community members’ GEMs based on their genome sequences [37], [38]. GEMs-FBA models are especially effective in revealing information about metabolic interactions underlying community assembly and function *in silico* [11], [29], [39]. Due to their mechanistic nature, in principle, GEMs-FBA models can be used to predict metabolic functions of any community with available GEMs in a defined environment without requiring any training data. However, their predictive performance is limited by a few factors. GEMs-FBA models are based on the assumption that each species adjusts their metabolic fluxes to optimize an objective function, typically related to maximizing biomass [37], which may not be relevant for certain conditions [40]. In addition, they do not explicitly account for intra-cellular regulation or non-growth mediated processes, such as toxicity arising from pH [41] or anti-microbials [23]. To evaluate GEMs-FBA models’ predictive capability on synthetic community data, we simulated the fluxes of fermentation end products and short-chain fatty acids (SCFA) using a synthetic community dataset [27]. This dataset consisted of combinations of 25 diverse human gut microbial species that are prevalent across individuals. Model simulations were performed using MICOM, which is an established method to perform GEMs-FBA model simulations for human gut microbial communities [29], [42] (**Methods**). The resulting Pearson correlations between predicted and measured community functions (i.e., SCFA production fluxes) range between -0.09 and 0.30 (**Figure 2**G). As a reference, a SAM that combines gLV dynamics to describe species abundance and a statistical model to map species abundance to butyrate production displays a Pearson correlation of 0.86 for butyrate [27]. These results show that GEMs-FBA models, which are primarily used to study the metabolic mechanisms shaping microbial communities, have limited predictive capabilities for the functions of synthetic microbial communities.

**Figure 2:**
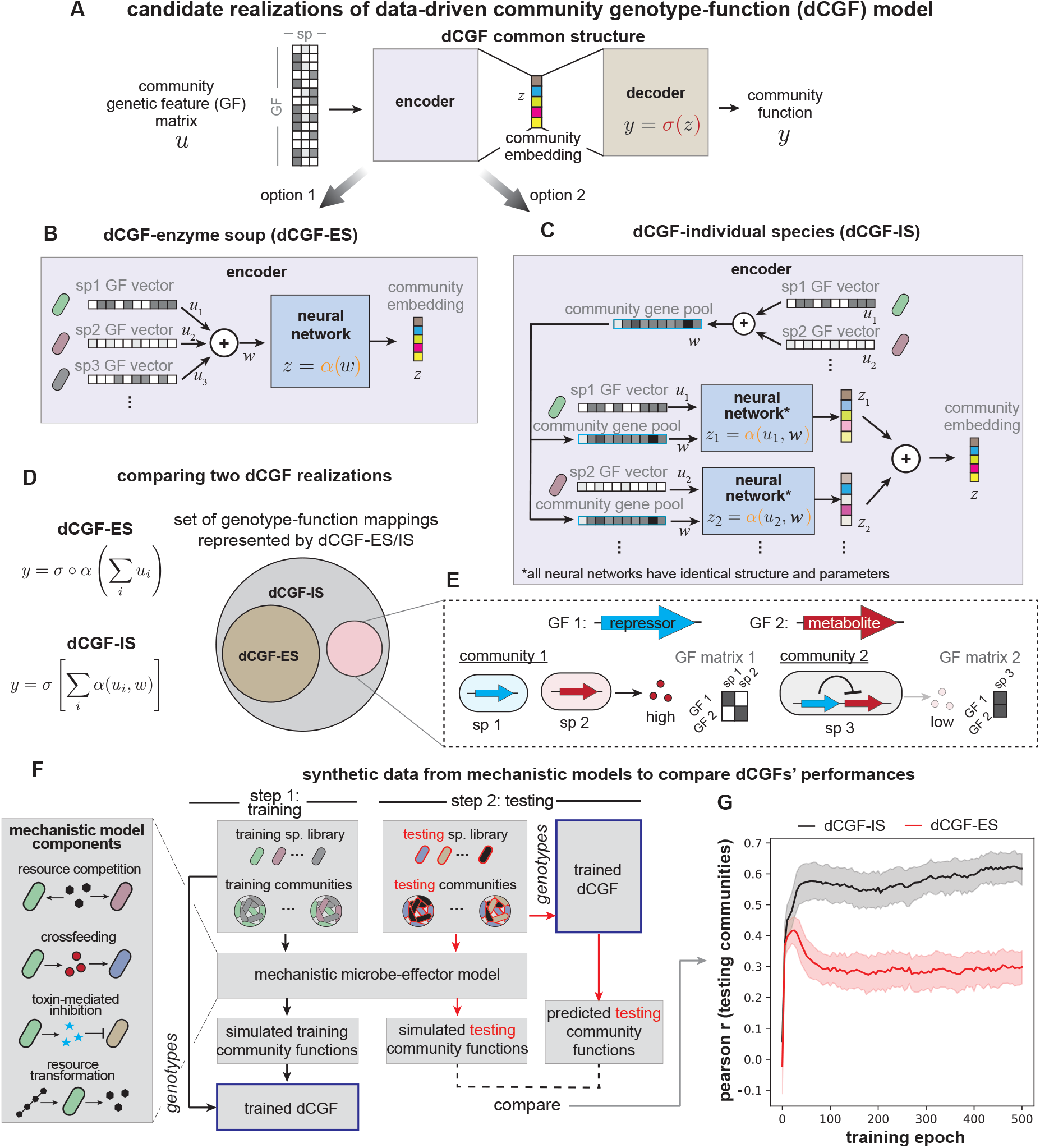
Candidate realizations of dCGF and their performances on synthetic data. (A) A common model structure of candidate dCGFs. The community GF matrix *u* is first fed into an encoder *z* = *α*(*u*) to extract the community embedding *z*. The community embedding *z* is then fed into the decoder *y* = *σ*(*z*), which is a fully connected neural network to predict community function(s). The two candidate realizations of dCGFs differ in their encoder structures. (B) The encoder of a dCGF-enzyme soup (dCGF-ES) model sums up the GF vectors of all species to form a community GF vector *w*, which is then fed into a fully connected neural network to output the community embedding *z*. (C) By contrast, the encoder of a dCGF-individual species (dCGF-IS) model process the genetic feature vector of each species individually. Specifically, the abundance-weighted GF vector of each species *i, u*_*i*_, and the community gene pool vector, *w*, are fed into a neural network to obtain each species’ functional embedding *z*_*i*_. The functional embeddings are summed up to become the community embedding *z*. The neural networks processing each species’ GF vector have identical weights and structures. (D) dCGF-IS is strictly more expressive than dCGF-ES. For every species genotype to community function mapping represented by dCGF-ES (buff-colored set), there exists a parametrization of dCGF-IS (gray-colored set) that can represent the same genotype-function mapping. A subset of community genotype-function mappings (pink-colored set in panel D) can be represented by dCGF-IS but cannot be represented by dCGF-ES. (E) An example of species genotype to community function mapping that cannot be represented by dCGF-ES. The two communities have different GF matrices, resulting in different output metabolite productions (high vs. low). However, dCGF-ES will make same predictions for the two communities since the column-sum of the two communities’ GF matrices are identical. (F) Mechanistic microbe-effector models (MEM) were used to generate synthetic data to train/test two dCGF realizations and compare their performances. Specifically, training and testing species were each assigned a random GF vector, which was then used to determine its MEM parameters, such as resource consumption rates and toxin production rates. Simulated function of MEMs from training communities were used to train dCGFs to predict the functions of testing communities. All species in testing communities do not appear in training data. The MEM model includes multiple modes of inter-species interactions and intra-cellular regulations. See **Methods** for more details on synthetic data generation. (F) Testing performances of the two different dCGF realizations for different training epochs. Testing performances were evaluated as the Pearson correlation between community functions simulated by ground-truth MEM and predicted by dCGF. Shaded areas quantify 95% confidence intervals arising from ten different random seeds used to generate the MEM parameters as well as to initialize neural network weights in dCGF (see **Methods**).

In sum, SAMs lack the ability to predict the functions of communities containing species that are absent from the initial training data. GEMs-FBA is currently the only widely used modeling framework to predict communities from species’ genome information and does not require training data. However, GEMs-FBA rely on rigid assumptions that are often difficult to verify, and they lack the flexibility to integrate high-throughput experimental data, limiting their predictive performances.

### Overview of dCGF: a data-driven model to predict community functions from its members’ genetic features

To address these gaps, we propose the data-driven community genotype function model (dCGF), as a proof-of-concept modeling framework to predict functions of synthetic microbial communities in a fixed environmental context from its members’ genetic features using tailored machine learning (ML) model structures. Compared to SAMs, dCGF lifts the representation of each species from a one-dimensional abundance value to a *n*_*g*_-dimensional genetic feature (GF) space, where typically *n*_*g*_ is much greater than 1. For example, *E. coli* K12 strain has over 4000 known genes and over 400 known metabolic pathways [43]. This enables prediction of communities assembled from at most 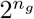 numbers of species with different combinations of GFs, including new species that are not present in training data. While dCGF takes genetic-level information as model inputs, in contrast to GEMs-FBA models, dCGF does not assume the communities optimize a specific objective function. Instead, the model has a tailored ML structure that has the flexibility to account for intra-cellular regulations and inter-species interactions, enabling it to integrate high-throughput experimental data.

Consider a synthetic microbial community composed of *n*_*s*_ species, where each species *i* is represented as a binary GF vector, indicating the presence/absence of *n*_*g*_ GFs harbored by species *i*. dCGF is a ML model that maps the GFs of all community members as well as their initial abundances, encoded as an *n*_*g*_ × *n*_*s*_ GF matrix *u*, to community function *y* (**Figure 1**H). Specifically, the *i*-th column of matrix *u* is the binary GF vector of species *i, v*_*i*_, scaled by its initial abundance *a*_*i*_: *u*_*i*_ = *a*_*i*_ · *v*_*i*_. The GFs can be chosen flexibly with varying granularity, depending on the availability of knowledge regarding the species’ genetic contents and their relevance to the community functions of interest. For example, some possible choices are the presence/absence of metabolic pathways or a subset of genes that are most related to the community function of interest based on expert knowledge. The output of dCGF is the emergent, quantifiable community function *y* of interest, ranging from its ability to degrade complex substrates, to inhibit a specific pathogen, or to produce a key metabolite. In principle, *y* can be a scalar, if one is interested in only one community function, or a vector, if multiple functions are measured/predicted simultaneously for each community.

In sum, a trained dCGF can predict functions of synthetic communities in a fixed environmental context that include new species, which are absent in training data and have known GFs (**Figure 1**I). However, the model’s predictive performance may be diminished if a given GF critical to a given community function is only present in that species and absent in all other species in the training set. Compared to SAMs, which cannot predict communities assembled from species beyond the training species, dCGF transforms this extrapolation task into interpolating the higher-dimensional GF space, a task where modern ML models, such as large neural networks, excel [44]. This data-driven modeling framework is particularly suitable for synthetic microbial communities. Compared to microbiome samples collected from natural environments, synthetic microbial communities can be assembled in defined environments using high-throughput lab automation techniques, thus providing rich experimental data for dCGF training. Further, by perturbing the presence/absence of the GFs of interest *in silico*, a trained dCGF can be used to discover the impact of GFs on a given community function (**Figure 1**I). This analysis can be used to guide the design of experiments that seek to uncover GFs that drive community functions partitioned across constituent community members.

### A biologically inspired model structure for mapping species genotype to community function

The mapping from species genotype to community function in dCGF may be captured by multiple ML model structures. To provide insights into the suitable model features and inductive biases for this task, we evaluate two models, referred to as dCGF-enzyme soup (dCGF-ES) and dCGF-individual species (dCGF-IS). dCGF-ES is a simple model that assumes community function can be determined by the total abundance of each GFs in the community. A similar assumption (i.e., enzyme soup) has been used in some GEMs-FBA models [37]. By contrast, dCGF-IS uses a structure inspired by the biological processes governing community assembly and function, including intra-species regulations and inter-species interactions.

At a high level, both candidate model structures belong to a broader class of neural networks called deep set [45]. Compared to other types of ML models, (i) the outputs of deep sets are invariant to input permutations and (ii) they can process inputs with flexible dimensions. Essential to modeling microbial communities, these two properties ensure that changes in species indexing (i.e., permuting columns of the community GF matrix) do not change dCGF predictions, and a trained dCGF can predict functions of communities of arbitrary size. In fact, while other ML models, such as convolutional neural networks (CNNs), can also take GF matrices as inputs, they do not possess such properties [45]. In particular, the sizes of input matrices for CNN must be fixed and therefore they cannot be applied to communities larger than the size of the largest training community. In addition, the output prediction using CNN changes with species indexing (i.e., column permutation).

The two dCGF candidates have a common encoder-decoder structure (**Figure 2**A). The encoder module extracts and compresses the information in the community GF matrix *u* to a lower dimensional community embedding *z*, which is then fed into a decoder module to output community function *y*: *y* = γ(*z*). Due to limited training data, we chose single layer neural networks to implement the decoders of both candidate models.

The two dCGF candidates differ in their encoder modules. dCGF-ES sums all columns of the community GF matrix (*u*), weighted by species’ initial abundances, as an input (*w*) to a fully-connected neural network (**Figure 2**B). The encoder structure in dCGF-ES uses a neural network to model the relationship between the sum of GFs and community embedding. This neglects the intra-cellular regulations and inter-species interactions that shape each species’ impact on the shared environment, and consequently community assembly and function.

The other candidate dCGF realization, dCGF-IS, employs a more sophisticated model structure inspired by the biological processes governing community assembly to map from *u* to *z* (**Figure 2**C). Specifically, in dCGF-IS, the community embedding *z* is determined by summing the contributions from individual species: *z* = ∑_*i* ∈*S*_ *z*_*i*_, where *S* is the set of all species and *z*_*i*_ represents the functional embedding of species *i*. This corresponds to the biological process where each species produces or consumes molecular effectors to influence their shared environment. The ability of each species to produce and consume the molecular effectors is affected by intra-cellular interactions and inter-species regulations. For example, the rate at which a species *i* produces a metabolite *y* may be determined by transcriptional regulators present in species *i* (e.g., GF 1) that activates or represses the metabolite producer (e.g., GF 2), which represents an intra-cellular regulation. Additionally, the presence of another metabolic pathway in species *j* (e.g., GF 3) may produce metabolic by-products, which can diffuse across cell membranes to reduce the rate of production of *y* by species *i*, which represents an inter-species interaction. We therefore assume that species functional Pembedding (*z*_*i*_) is jointly determined by its own GFs (*u*_*i*_) and the community gene pool (*w* = ∑_*i*_ *u*_*i*_) through intra-cellular regulations and inter-species interactions, respectively. We applied a neural network *z*_*i*_ = *α*(*u*_*i*_, *w*) to capture these potentially highly complex regulations and interactions.

The mathematical function of the encoder *α*(·, ·) is assumed to be identical across all species and does not change for different species. This assumption implies that there is a universal, learnable rule to map species’ GFs to their contributions to community function, laying the foundation to reuse this mapping (*α*) when predicting the impact of new species with known GFs on a community function. A similar assumption has been used previously [36], where the authors assumed there is a universal linear function, that maps each species’ presence/absence of genes to their resource consumption/production capabilities in consumer resource models. While the same linear function was applied to all species, the differences in the presence/absence of genes in each species yields different resource consumption rate constants among species. In comparison, dCGF-IS replaces the linear function with a more flexible neural network in the encoder. Therefore, using a universal mathematical form *α*(·, ·) for the encoder of every species does not mean that all species contribute to community embedding *z* identically. In fact, species have different GFs (*u*_1_ ≠ *u*_2_), hence their functional embeddings are different (i.e., *z*_1_ = *α*(*u*_1_, *w*) ≠ *α*(*u*_2_, *w*) = *z*_2_). Because a universal encoder is applied to all species, from a computational complexity point of view, dCGF-IS and dCGF-ES both have 𝒪(*n*_*g*_) parameters regardless of the number of species.

To compare the two candidates’ model capacity, we show analytically that dCGF-IS is strictly more expressive than dCGF-ES (**Supplementary Note 1**). This means that for every genotype-function mapping represented by dCGF-ES, there exists a parameterization of the neural networks in dCGF-IS that can produce the same mapping. Conversely, certain genotype-function mappings created by dCGF-IS cannot be produced by dCGF-ES, regardless of how the neural network parameters are chosen (**Figure 2**D, **Supplementary Note 1**). This is particularly relevant in the presence of intra-cellular regulations and inter-species interactions. In particular, for two communities with different GF matrices but identical GF matrix column sums (i.e., ∑_*i*_ *u*_*i*_), dCGF-IS can be trained to output different community functions, while dCGF-ES cannot (pink subset in **Figure 2**D). These differential community functions may arise from intra-cellular regulations and inter-species interactions. For instance, community 1 and 2 in **Figure 2**E both have a repressor gene and a metabolite production gene (i.e., identical GF matrix column sums). However, since the two genes are distributed in different (the same) species in community 1 (community 2), metabolite production is high (low) due to the absence (presence) of intra-cellular regulation. Therefore, dCGF-IS has more capacity to account for intra-cellular regulations and inter-species interactions than dCGF-ES.

To evaluate the effects of intra-cellular regulation an inter-species interactions on model prediction performance, we generated synthetic data using mechanistic microbe-effector ordinary differential equation models (**Figure 2**E). This approach enables a comparison of dCGF-ES and dCGF-IS performances with a known ground truth model, noiseless measurements, and abundant training data. The mechanistic model explicitly accounts for how each species’ GFs yields different production and consumption rates of a half dozen molecular effectors, including metabolites that are cross-fed, resources that are consumed, transformed resources (e.g., polysaccharide breakdown products produced from fiber), and toxins [14], [23], [26]. We used mathematical functions of various forms to map species’ GF vectors to parameters in the microbe-effector ODE models, such as resource consumption and toxin production rate constants (see **Methods** for ground-truth models). We trained dCGF-IS and dCGF-ES using the same methods (**Methods**) on the same synthetic dataset, which were simulated communities assembled from a set of species, which we call training species. The trained candidate models were used to predict communities assembled from a set of species with different presence/absence of GFs, which we call testing species (**Figure 2**E). This is a challenging prediction task since the testing species do not appear in the training data. Our numerical simulation shows that while both candidate realizations can learn species genotype to community function mapping from data, dCGF-IS reduces fitting residuals in training data (**Supplementary Figure 1**) and displays a substantial improvement in prediction performance for communities assembled entirely from testing species (**Figure 2**F).

In sum, dCGF-IS is more effective in capturing species genotype to community function mappings in microbial communities, whose functions of interests are shaped by intra-cellular regulations and inter-species interactions. Thus, in all subsequent analysis, we will apply dCGF-IS to experimental datasets.

### dCGF accurately predicts functions of communities assembled from training species

Compared to the dataset created *in silico* in **Figure 2**, experimental synthetic microbial community datasets *in vitro* are often limited in sample size and corrupted by substantial measurement noise and noise arising from the stochastic nature of molecular reactions. We evaluated the predictive capability of dCGF-IS using published experimental synthetic community datasets [27], [36], [46], [47] (**Figure 3**A). Each of these synthetic community datasets contains quantitative measurements of combinations of species in a defined and controlled environment. Hence, these data have offered mechanistic insights into microbial community assembly and function with applications ranging from precision probiotics that decolonize pathogens to restoration of gut homeostasis by production of gut health-promoting metabolites. The datasets we use include synthetic microbial communities assembled from diverse species *in vitro*, performing tasks ranging from production of health-relevant metabolites [27] in synthetic human gut communities (acetate, butyrate, lactate, and succinate datasets, 576 communities assembled from 25 species), to inhibition of the human gut pathogen *Clostridioides difficile* (*C. difficile*) by commensal human gut bacteria [46] (*C. difficile* inhibition dataset, 180 conditions assembled from 14 species), to denitrification in synthetic soil microbial communities [36] (nitrate and nitrite datasets, 186 communities assembled from 12 species), to degradation of complex fibers by soil microbes [47] (starch degradation dataset, 53 communities assembled from 6 species).

**Figure 3:**
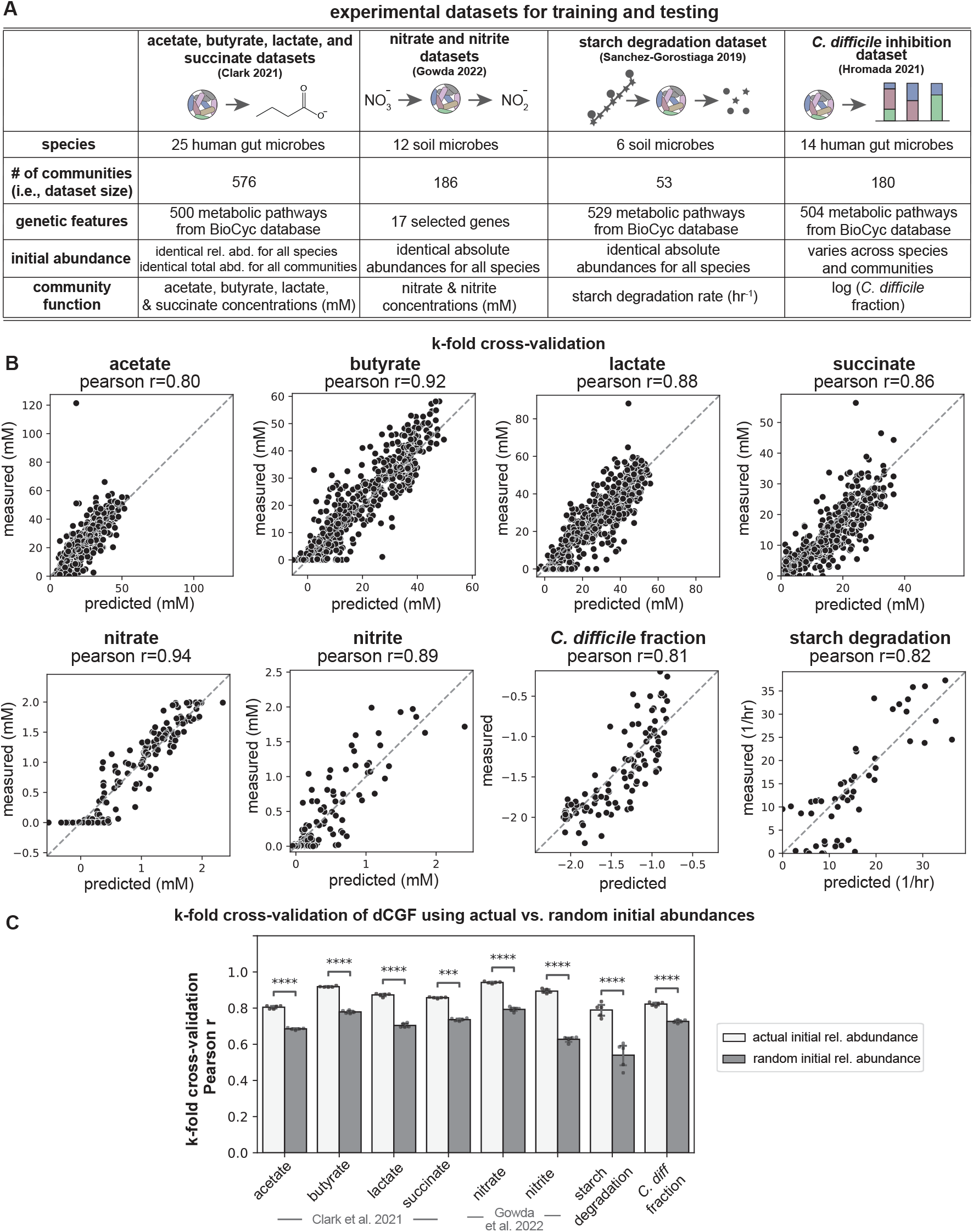
Evaluation of dCGF performance on experimental data using *k*-fold cross-validation. (A) A summary of the published experimental datasets [27], [36], [46], [47] used to train and test dCGF-IS. (B) Scatter plots of dCGF predicted and measured community functions for all experimental datasets. Five different random seeds were used to split data into *k* folds and to initialize neural network parameters. Predicted function of a community is the average of the five predictions. (See **Methods**.) (C) Bar plot of *k*-fold cross-validation performances of dCGF-IS using actual vs. random initial species relative abundances. Error bars represent 95% confidence intervals arising from five different random seeds to initialize neural network parameters. ***: *p <* 10^−3^, ****: *p <* 10^−4^ using independent t-tests.

The species’ GFs can be encoded in many ways utilizing different types of genetic information with different granularities, ranging from the presence/absence of genes to metabolic pathways. We determined the GFs for these datasets based on their relevance to the community function of interest. Based on prior knowledge, we assume that a community’s ability to produce acetate, butyrate, lactate, succinate, to inhibit *C. difficile*, and to degrade starch are strongly driven by metabolic activities [48], [49]. We therefore used one-hot encoding of each species’ presence/absence of metabolic pathways, as curated in the BioCyc collection of Pathway/Genome Databases (PGDBs) [50], to represent their GF vectors. BioCyc is a regularly maintained collection of databases that contains PGDBs for over 20,000 organisms, where each PGDB stores information of metabolic pathways, reactions, enzymes, genes, operons, transcription factors, and DNA binding sites present in the genome of a species. The presence/absence of these GFs in each species can be found directly in BioCyc. Metabolic pathways and reactions in BioCyc are widely used to construct GEMs for FBA [51]. Hence, it is an accessible, context-agnostic, and reliable source to generate GF-vectors for metabolite-related community functions. Lower granularity genetic information (i.e., metabolic pathways) was used to encode species’ GFs in these datasets, as opposed to higher granularity ones (e.g., genes), based on training data availability considerations. Specifically, the sample sizes of these experimental datasets range between 53 to 576, which are much smaller than the thousands of genes present in each species [43]. For the remaining nitrate and nitrite datasets, 17 genes have been identified based on expert knowledge as highly relevant to denitrification [36], and hence their presence/absence were one-hot encoded as species’ GFs.

We evaluated the prediction performance of dCGF-IS on all experimental datasets using *k*-fold cross-validation by training the model on a set of communities and evaluating the model’s predictive capability on a different set of communities. This is the standard performance metric for data-driven SAMs, where each species may appear in both training and testing data. We trained one dCGF for each dataset (i.e., each community function), and the resultant Pearson correlation coefficients were between 0.80 and 0.94 (median 0.87) for all 8 datasets (**Figure 3**B). Data-driven SAMs have previously been applied to the acetate, butyrate, lactate, and succinate datasets, resulting in similar *k*-fold cross-validation performances [28], [31]. Prediction performance of dCGF increased with the amount of available data used for training (**Supplementary Figure 2**). Performance also generally increased with community size for most datasets (**Supplementary Figure 3**), which is consistent with previous observations from GEMs-FBA models [29]. dCGFs’ *k*-fold cross-validation performances were largely insensitive to the number of nodes in the neural networks (**Supplementary Figure 4**) and were consistent for a large range of training epochs (**Supplementary Figure 5**). This observation is consistent with the robustness in prediction performance of high capacity ML models, such as deep neural networks, to variation in hyperparameters [44]. Therefore, when the testing communities are assembled from species already present in training data, dCGF-IS has comparable prediction performances to the state-of-the-art data-driven SAMs.

As the input to dCGF-IS, each column in the community GF matrix, representing a given species, is scaled by its initial relative abundance. This accounts for variation in the potential impact of a given species on community function due to changes in its initial abundance [33]. To evaluate whether incorporating species’ initial abundance improves dCGF performance, we randomly perturbed the species’ initial abundances and compared the *k*-fold cross-validation performances. Using the same training/testing procedure and hyper-parameters, the *k*-fold cross-validation performance using the actual initial abundances was substantially higher than random initial abundances (**Figure 3**C, **Supplementary Figure 6**). Hence, scaling the input of dCGF-IS based on initial species abundances is essential to its performance.

By perturbing the community GF matrix *in silico*, a trained dCGF-IS can simulate the extent to which each GF in each species impacts community function to generate hypothesis (**Figure 4**A). We performed sensitivity analysis on dCGF-IS trained with the butyrate dataset, where the relevant biomolecular mechanisms are well-studied [52]. Three of the top 5% metabolic pathways that enhance butyrate production were related to lipoate biosynthesis and incorporation (**Figure 4**B). Lipoate is an essential co-factor required for pyruvate dehydrogenase [53], [54], which is the primary enzyme for acetyl-CoA biosynthesis [55]. Acetyl-CoA is the prevalent substrate for butyrate synthesis [52]. In addition to lipoate-related pathways, acetyl-CoA fermentation to butyrate [15], [52], sulfate reduction [27], and inositol degradation pathways [56], [57], which are known to affect butyrate production and/or butyrate producer growth in the literature, have sensitivity magnitudes in the top 5%. We computed sensitivities for the acetate, lactate, and succinate datasets in **Supplementary Figure 7**. The metabolic pathways with the largest sensitivity magnitudes for acetate and lactate were similar to those of butyrate, while many of those for succinate were different. This could be explained by the strong correlation between acetate, butyrate, and lactate in experimental datasets as lactate and acetate can be converted into butyrate (**Supplementary Figure 7**, insets). These results imply that sensitivity analysis of dCGF-IS can guide hypotheses to test the predicted mechanisms experimentally or to guide selection of genetic targets for further investigation.

**Figure 4:**
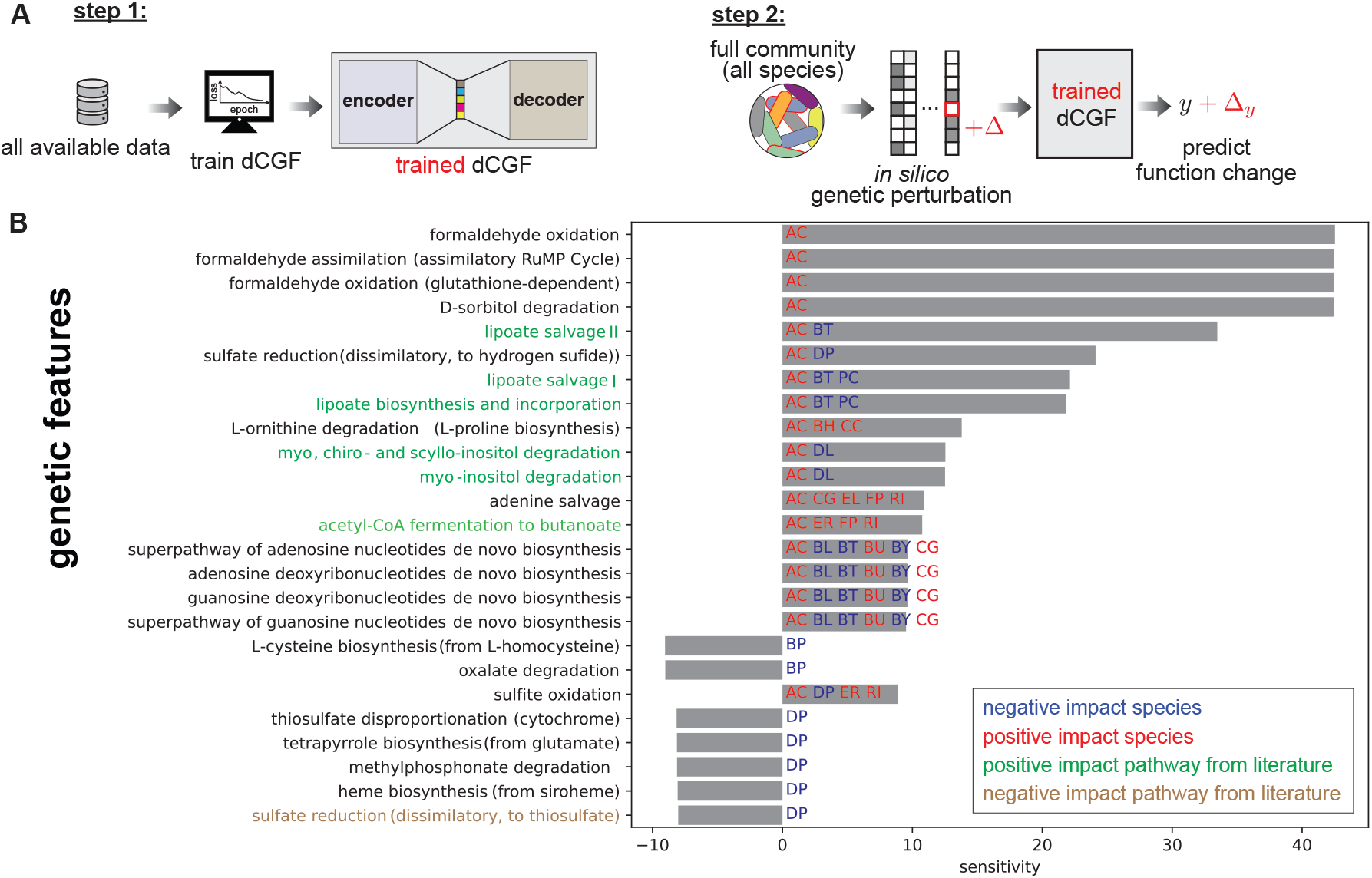
Sensitivity analysis of dCGF to evaluate the impact of each GF on community function. (A) Workflow to compute sensitivities of each GF. All available data for a community function was first used to train a dCGF-IS. GF matrix of a full community that includes all possible community members was used as the nominal input to the trained dCGF-IS. To perturb the nominal GF matrix, each GF was turned on (off, respectively) if it was off (on, respectively) in the nominal GF matrix. The perturbed community function was then predicted using the trained dCGF-IS and the perturbed GF matrix. The difference between nominal community function and perturbed community function was used to evaluate sensitivity of a GF in a species. Finally, the sensitivity of a GF is computed by averaging across all species and all random seeds (not drawn in figure). See **Methods** for more details. (B) Sensitivities of GFs with the top 5 % magnitudes computed from dCGF-IS for the butyrate dataset. Species where the GFs are present are labeled on top of each GF. Species that were found to positively (negatively, respectively) impact community butyrate production on average were labeled red (blue, respectively). The impact of a species on community butyrate production is the difference between the average butyrate in communities with and without this species. GFs that are known to positively (negatively) impact microbial butyrate production are shown in green (brown, respectively).

In sum, dCGF-IS can predict functions of communities assembled from training species in experimental datasets, and sensitivity analysis based on trained dCGF-IS can generate hypotheses about the impacts of GFs on community function.

### dCGF accurately predicts functions of communities involving new species absent from training data

A major advantage of dCGF over SAMs is the ability to predict functions of communities composed in part or entirely of new species that are absent in training data. For example, this could be used to inform the selection of species for building synthetic communities from the bottom-up to study and optimize community functions of interest using an expanded design-test-learn cycle paradigm [31]. We therefore evaluated the prediction performance of a model trained on communities assembled from a set of training species excluding one community member (i.e., “testing species”). We then used the model to predict functions of all communities initially including this member (“leave-one-out” test) (**Figure 5**A). Hence, each testing community always includes one species that has never been used for model training. For a dataset containing communities assembled from *n*_*s*_ species, we labeled each species as testing species once, leading to *n*_*s*_ leave-one-out test scores (**Figure 5**B). The model achieved overall high prediction performance across all 143 combinations of community functions and testing species (median Pearson correlation 0.85) and 95% of the combinations displayed a Pearson correlation greater than 0.5 (**Supplementary Table 2**). For the acetate, butyrate, lactate, and succinate datasets, dCGF-IS’ prediction performances were substantially higher than those of GEMs-FBA models (**Figure 1**G). These data demonstrate that dCGF-IS can achieve substantially higher prediction performance than the current mainstream modeling method to predict functions of communities including new species. While GEMs-FBA models are knowledge-based and dCGF is data-driven, our results highlight that high-throughput synthetic community experimental data can be leveraged to infer the connection between species’ genotypes and community function. This, in turn, enhances prediction performance—an area for which GEMs-FBA models are not designed.

**Figure 5:**
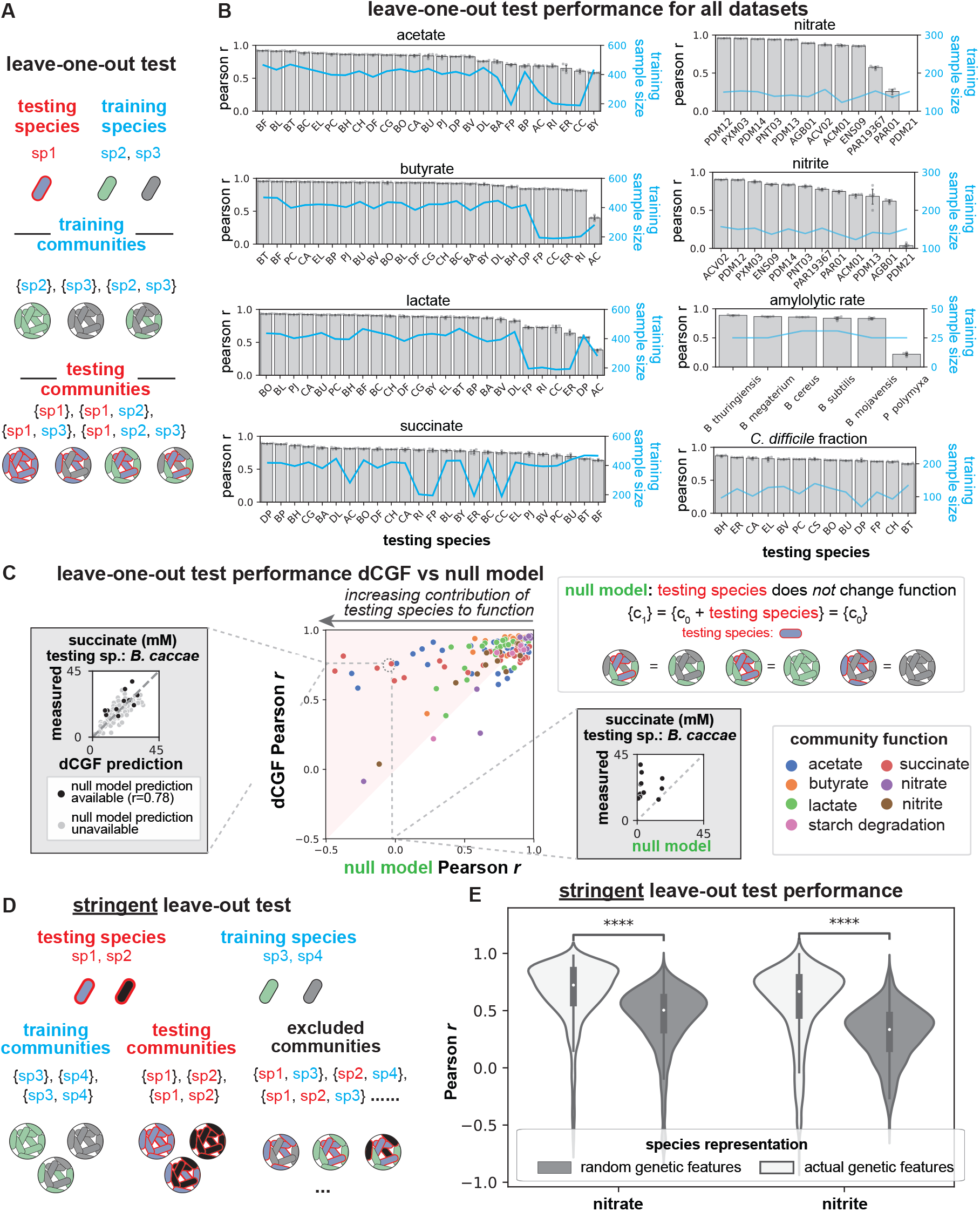
Performance of dCGF-IS to predict functions of communities including new species with known GFs that are absent in training data. (A) Schematic of the “leave-one-out” test. One species is labeled as testing species and all communities initially including the testing species are used as testing data. The rest of the communities in the dataset are used for model training. (B) Leave-one-out performances for all datasets and all possible testing species. Bar plots show mean Pearson correlation between model prediction and measured communities functions for the testing communities. Error bars indicate 95% confidence intervals arising from five random seeds to initialize dCGF parameters. Blue solid lines show training sample size (i.e., number of communities used for training). (C) Comparison of leave-one-out performance between dCGF and a null model for all datasets and all possible testing species. Upper right box shows examples of the null model, which makes predictions assuming the testing species does not change community function. Each dot in the scatter plot shows the leave-one-out performance of dCGF and null model for one dataset using one species as left-out. Colors of the dot represent different datasets (i.e., community functions). Red shaded region are dataset/testing species combinations where dCGF outperformed the null model. Gray inset boxes show one example of predicted vs. measured community function. (D) Schematic of the “stringent leave-out” test. For each dataset, a set of *n*_*l*_ species were labeled as testing species and the rest of the species were labeled as training species. Communities composed entirely of the testing (training, respectively) species were used for testing (training, respectively). Communities including both testing and training species were excluded. (E) Stringent leave-out performances for nitrate and nitrite datasets with all possible combinations of four testing species. Species were encoded either using their actual GFs (dark gray violin plots) or random GFs (light gray violin plots). ****: *p <* 10^−4^ using one-sided Mann-Whitney U test.

The observed variation in leave-one-out test scores across species in a given dataset may arise from different training sample sizes. The microbial communities in the published datasets were assembled in a biased way such that certain species were more or less frequent in the communities. Therefore, the training data sample size changes when different species were labeled as testing species (**Figure 5**B). For example, the butyrate producers *A. caccae* (AC), *C. comes* (CC), *E. rectale* (ER), *F. prausnitzii* (FR), and *R. intestinalis* (RI) are present in 51%, 67%, 66%, 66%, and 65% communities, respectively, in the acetate, butyrate, lactate, and succinate datasets. Hence, the available training data size was low when they were used as testing species. For acetate, butyrate, and lactate datasets (**Supplementary Figure 8**), testing species with low training sample sizes displayed moderate leave-one-out test scores. Hence, leave-one-out test scores for these testing species-dataset combinations could potentially improve given more training data. By contrast, for nitrate, nitrite, starch degradation, and succinate datasets, there were no clear correlations between training sample size and leave-one-out test scores (**Supplementary Figure 8**). Specifically, when used as training species, PDM21 in nitrate and nitrite datasets and *P. polymyxa* in starch degradation datasets have poor leave-one-out test scores while their corresponding training sample sizes are similar to other species’ in the dataset (**Supplementary Figure 8**). In these cases, the reduced prediction performances may be due to unique GFs in the testing species that are not present in any training species or unannotated GFs. Additionally, the testing species may harbor unique combinations of GFs, which may interact to give rise to intra-cellular regulations and inter-species interactions that cannot be learned from training data.

A high leave-one-out prediction performance may stem from the testing species not sufficiently changing the community function. For example, suppose x is the testing species, the function of a community c_0_ may be similar to that of community c_1_ = c_0_ + x, when the addition of x does not directly change community function (e.g., the species could be excluded from the community over time). This makes it possible to predict c_1_ function using a hypothetical “null model”, where the function of c_0_ is used as a predictor for the function of c_1_ (**Figure 5**C, upper right box). To evaluate this possibility, we compared the performances of the null model to dCGF-IS for all possible testing species-dataset combinations. dCGF-IS outperformed the null model in 66% of the combinations, including many hard to predict combinations where the null model failed (**Figure 5**C, red shaded region in scatter plot). Indeed, out of all 24 combinations where the null model displayed a Pearson correlation less than 0.5, dCGF-IS only slightly underperformed in one combination. By contrast, the combinations where the null model outperformed dCGF were mostly composed of easy to predict combinations where both models performed well, where the Pearson correlations were greater than 0.5 (**Figure 5**C, scatter plot uncolored region). In these cases, the testing species may not substantially contribute to the given community function. Furthermore, the overall improvement of dCGF-IS to the null model is statistically significant (**Supplementary Figure 9**). These results further demonstrate that dCGF-IS extracted useful information from the species’ GFs to predict community functions.

Ideally, a trained dCGF-IS can predict communities composed entirely of new species with known GFs, as we showed using the synthetic dataset in **Figure 2**E-F. We call such performance tests “stringent leave-out” tests, where communities assembled from a set of training species sp_1_, …, sp_*n*_ are used to train a dCGF to predict functions of communities assembled from a set of new testing species sp_*n*+1_, …, sp_*m*_ (**Figure 5**D). When applying this approach, any community that includes both training and testing species will be discarded (**Figure 5**D), limiting the availability of training/testing data. The published experimental datasets were not designed for stringent leave-out tests. For instance, in the starch degradation and *C. difficile* inhibition datasets, there does not exist any combination of four testing species such that the resulting training sample size is greater than 10. In this case, the training data sample size is insufficient to train a model with hundreds of input GFs.

However, for the nitrate and nitrite datasets, each species is encoded by only 17 manually curated GFs (i.e., smaller input dimension), allowing dCGF-IS to be effectively trained with smaller training sample size. We therefore performed stringent leave-out tests on these datasets, labeling different 4-species combinations as testing species. Overall, partition of training and testing species led to a median training sample size of 60, whereas the training sample sizes for leave-one-out tests on the nitrate and nitrite datasets were larger than 120. For comparison, we also performed stringent leave-out test by encoding all species in the nitrate and nitrite datasets using random GF vectors, instead of the actual presence/absence of GF in these species’ genomes. dCGF-IS trained using randomly sampled GFs had Pearson correlation medians of 0.56 and 0.23, respectively, for the nitrate and nitrite datasets. By contrast, the resultant stringent leave-out tests had Pearson correlation medians of 0.72 and 0.67, respectively, when dCGFs were trained using species’ actual GFs (**Figure 5**E). The substantially higher stringent leave-out scores implies that dCGF learned useful information about the mapping between species genotypes and community function mapping from synthetic community data.

For the acetate, butyrate, lacate, and succinate datasets, regardless of the partition of training/testing species, the training sample size is much smaller than the number of GFs (**Supplementary Figure 10**). As a result, dCGF-IS failed the stringent leave-out test for acetate, butyrate, and lactate datasets using experimental data only (**Supplementary Figure 10**). To address the lack of training data, we incorporated biological knowledge on the production and consumption of these metabolites into model training. This is a widely used practice in physics-informed ML [58]. Specifically, we applied a data-augmentation approach in physics-informed ML [58], [59], where we provided additional synthetic data for dCGF training (**Methods, Supplementary Figure 10**) to induce observational biases to improve model prediction performance. The synthetic data we provided highlighted the impact of three driver metabolic pathways (i.e., GFs) to butyrate, acetate, and lactate production and consumption, including CoA transferase pathway, butyrate kinase pathway, and the lactate to pyruvate fermentation pathway. This was accomplished by first creating synthetic species with random GFs, and then assigning low/high butyrate, acetate, and lactate values produced by these species based on the presence/absence of the three driver pathways. Because dCGF is overparameterized, when it was pre-trained using these synthetic data, its neural network parameters biased towards a solution that put heavier weights on the three driver pathways. Using these biased parameters to initialize dCGF training when given experimental data resulted in substantially improved stringent leave-out performances, achieving Pearson correlations 0.61, 0.71, and 0.51 for acetate, butyrate, and lactate, respectively (**Supplementary Figure 10**). Sensitivity analysis of the pre-trained model also revealed the three driver metabolic pathways among the most important pathways for butyrate production (**Supplementary Figure 11**). These numerical experiments showcase that dCGF-IS can be integrated with mechanistic knowledge of community assembly and function to improve its prediction performance.

In sum, using leave-one-out and stringent leave-out tests, we demonstrated that, given sufficient and properly collected training data, dCGF-IS could be a promising data-driven approach to predict communities including new species with known genetic information. Hence, dCGF provides a new data-driven approach to model microbial community functions based on genetic information.

### dCGF provides a data-driven approach to quantify and compare species’ functional roles

Quantifying and comparing the effects of microbes on a given community function can provide deeper insights into their functional roles on a given community function [60], [61]. Such insights may uncover potential keystone species or design synthetic communities that optimize a function of interest [31]. In the absence of functional data, a widely used approach is to directly compare the species’ genetic contents using a phylogenetic tree based on genetic information. However, this approach neglects interdependencies among GFs arising from intra-cellular regulations, or differential “weights” of GFs to the function of interest [60]. For example, consider a wild-type strain sp_1_ and its mutant sp_2_, which lacks a polysaccharide utilization loci (PUL) to degrade a fiber y. While the genetic contents of sp_1_ and sp_2_ are largely similar, making them close to each other in a phylogenetic tree, their fiber degradation capabilities are very different. In this case, comparing species’ genetic contents alone cannot identify GF with differential weights to quantify their functional similarities.

dCGF-IS provides a data-driven approach to deal with these complexities by extracting and compressing information from species’ GF vectors and community function data. Specifically, the encoder of dCGF-IS automatically transforms the higher dimensional GF vector (*u*_*i*_) of each species *i* to a low dimensional species’ functional embedding (*z*_*i*_). This embedding represents the contribution of species *i* to community state (i.e., community embedding) *z*, which is directly related to a particular community function. To demonstrate dCGF’s capability to extract species’ functional contributions from genotypes and perform functional comparison, we trained a dCGF-IS using all available data for each experimental dataset in **Figure 3**A. For each community function of interest, we then fed the GF vectors representing each species *i* with unity abundance (*v*_*i*_) to the encoders of the trained dCGFs, which output each species’ functional embedding (**Figure 6**A). We used unity abundance to eliminate the impact of abundance on species functional comparison. The functional embeddings *z*_*i*_ summarize the multidimensional impact of species *i*’ on a particular community function, which is learned from data to account for the interdependencies among GFs as well as their differential weights on the function of interest. To compare species’ functional roles, we performed principal component analysis (PCA) directly using species’ GF vectors (**Figure 6**B) or their functional embeddings computed by dCGFs (**Figure 6**C). For all datasets, the scores of the first principal component (PC 1) calculated using species’ functional embeddings were strongly correlated with their experimental functional contributions (**Figure 6** D), which was calculated as the mean changes in the community functions with and without that species in all available experimental data. By contrast, these strong correlations disappeared when dCGF was trained using species encoded by random GF vectors (**Supplementary Figure 12**). In sum, these observations indicate that the encoder of a trained dCGF-IS can extract meaningful functional representations from the GF vectors of the species.

**Figure 6:**
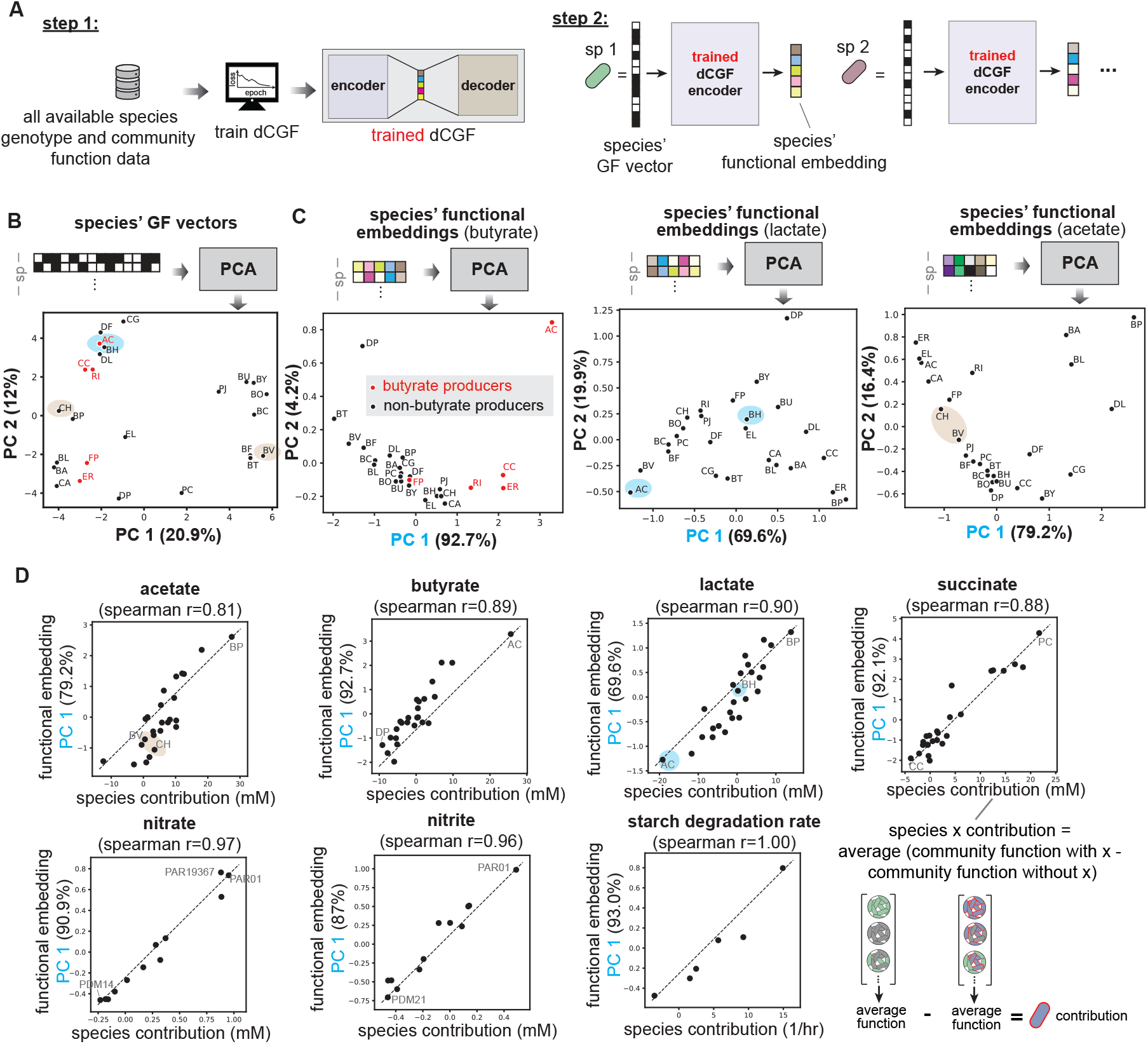
Species functional embeddings are highly relevant to their functional contributions in the communities. (A) Species’ functional embeddings were derived by first training a dCGF using the entire dataset. The GF vector of each species (with unity abundance) is then fed into the trained dCGF encoder to obtain its functional embedding. (B)-(C) Principal component analysis (PCA) of the species’ GF vectors (B) and functional embeddings (C) for butyrate, lacate, and acetate datasets. (D) Correlation between species’ scores for PC 1, computed using their functional embeddings, and their contributions to each community functions. The contribution of species x to a community function is computed as the difference between the average measured functions of communities including x and the average measured functions of communities excluding x in the entire dataset.

As a confirmatory result, *A. caccae* (AC), *C. comes* (CC), *E. rectale* (ER), *R. intestinalis* (RI), and *F. prausnitzii* (FP) produce butyrate in monoculture and were known as butyrate producers. The butyrate producers display high PC1 scores according to PCA of species’ functional embeddings (**Figure 6**C, left), which may be regarded as a direct measure of species’ butyrate producing capabilities. By contrast, PCA of species’ GFs did not show such a trend, with the butyrate producers not showing high (or low) scores for the first two PCs (**Figure 6**B). Further, comparing **Figure 6B** and C, the first principal components computed using species’ functional embeddings explain much more variability (between 69.6% and 92.7%) than the first principal components computed using GFs directly (20.9%). These observations show that the dCGF-IS encoder process each species’ GF vector to extract functional embeddings that are closely related to their contributions to community functions.

While PCA computed using species’ GF vectors (**Figure 6**B) did not depend on the function of interest, the one computed using species’ functional embeddings differed for different functions of interest (**Figure 6**C). This underscores the crucial, yet often overlooked, fact that functional “similarity” between species must be defined in the context of a specific function For instance, from a genotype perspective, *A. caccae* (AC) and *B. hydrogenotrophica* (BH) were close according to PCA of their GFs (blue shade in **Figure 6**B) as well as according to the phylogenetic tree of all species in the lactate dataset [27]. However, their contributions to community lactate production were dramatically different according to PCA of their functional embeddings (blue shade in **Figure 6**C middle panel) or according to their experimentally measured functional contributions [28] (see also **Figure 6**D, lactate panel). Conversely, *C. hiranonis* (CH) and *B. vulgatus* (BV) were far apart from a genotype perspective (brown shades in **Figure 6**B) as well as according to the phylogenetic tree of all species in the acetate dataset [27]. However, these species were similar in community acetate production and consumption (brown shade **Figure 6**C right panel and **Figure 6**D, acetate panel). In sum, functional embeddings derived from dCGF can quantify the contributions of species towards a given community function. They are more effective than GF vectors to quantify functional similarity among species (**Figure 6**B-D), which can be leveraged to design communities to optimize a function of interest. Notably, for all community functions studied, the strong correlation between species’ functional embeddings and functional experiment data indicates that the encoder of dCGF-IS learns from experimental data to compress and extract genetic information to obtain each species’ functional contributions.

## DISCUSSIONS

Data-driven models that learn species genotype to community function mappings from high-throughput synthetic microbial community data have the promise to substantially accelerate synthetic community design and investigation. The selection of species for constructing synthetic microbiomes is a key factor in determining the types of discoveries that can be made, yet we currently lack a systematic framework to guide this process. The dCGF modeling framework enables the selection of species/strains to optimize a given function or fill knowledge gaps in data-driven models, enabling an expanded design-test-learn framework [31]. These models provide a major advance in the prediction of functions of communities that include new species with known GFs absent from training data. Using explainable ML tools, they can also be applied to quantify the impact of each GF on community function *in silico*. These capabilities are absent in SAMs. While mechanistic GEMs-FBA models have similar capabilities, they are difficult to integrate with high-throughput experimental data, thus limiting predictive performances.

In the broader context of predicting microbial community functions from species’ genotypes, dCGF is designed specifically for synthetic microbial communities in fixed environmental contexts as opposed to natural microbiomes. Synthetic communities are composed of designer-specified species with known abundances, making it possible to use their GFs and initial abundances as model inputs. By contrast, compositions of natural microbiome samples must be first inferred from metagenomic or 16S rRNA gene sequencing data, presenting an additional layer of challenges to the data-driven prediction of natural microbiomes [29], [62]–[64].

Datasets for synthetic microbial communities are collected from well-defined lab environments with controlled variability across samples. Therefore, dCGF can be applied to quantitatively predict functions of new communities assembled in the same environment as training data. By contrast, natural microbiome datasets are collected from distinct environments that have uncontrolled and often unknown sources of variability. Due to these challenges, existing ML models for natural microbiome data has primarily focused on classification problems (e.g., disease vs healthy), instead of quantitative prediction. Quantitative prediction can be critical for to designing communities to optimize target functions such as metabolite production and fiber degradation [27], [47]. Furthermore, synthetic microbial communities are typically composed of no more than 100 species due to limitations in highly parallelized culturing [27], [46], [65]. Hence, the types of species in a synthetic community will likely need to be augmented with new species or strains to further optimize a target function and discover genetic features that impact such function. By contrast, natural microbiomes can contain thousands of species [66] and many public metagenomic datasets are available for the same phenotypes (e.g., metagenomic datasets relating to colorectal cancer). These make it feasible to train one classification model by aggregating different datasets [63], which can contain several thousands of species in training data. Consequently, the presence of unknown species absent from training data is uncommon and often neglected in natural microbiome datasets. Overall, dCGF is tailored and most suitable to quantitatively predict community functions from defined microbial communities in defined environments. Addressing the major challenges facing ML for natural microbiome data will require other model structures together with advances to process metagenomic data, including generating metagenome assembled genomes.

Similar to other ML models, the prediction and interpretation performance of dCGF may benefit from GF selection and pre-processing [67]. This is evidenced by the high accuracy and low variability of the model’s performance for the nitrate and nitrite datasets (**Figure 3, Figure 5**), where the genes most relevant to the community functions are manually curated as GFs. Such a manual GF selection strategy may be applicable to communities where the critical GFs responsible for a given function are known *a priori*. For example, synthetic communities assembled from engineered strains often contain inter-species interactions mediated through engineered quorum sensing, toxin production, and toxin protection genes [23]. The presence/absence of these genes can be encoded in their GF vectors. In addition, transcriptional profiling data could be used to identify and then remove GFs that are known to be silent in a given environment. Furthermore, additional relevant features, such as the presence/absence of PULs related to a media fiber component that are induced in the specific environment, could be included. Finally, statistical tools, such as independent component analysis [68], and unsupervised ML tools, such as variational auto-encoders, could be used for preprocessing of the GFs to reduce the dimension of the input space to improve model prediction performance while often sacrificing interpretability.

In addition to genetic information, the inputs to dCGF can be appended with vectors representing environmental contexts, such as concentrations of different medium components (e.g., concentrations of various amino acids and sugars). This extended form of dCGF can be trained on synthetic microbial communities assembled in different environmental contexts to predict community functions in new environmental contexts, which may facilitate community design to enhance their functions [69]. For example, by appending concentrations of different types of sugars and amino acids to the GF vectors as inputs to dCGF, functions of communities measured from different defined media may be leveraged to infer functions of communities assembled in a new medium with different sugar and amino acid concentrations. These extended dCGF models could reveal the effects of environmental contexts on community functions and uncover the complex interplay between genetic information, environmental contexts, and community function to facilitate community design and optimization. Finally, a training dataset with the proper genetic variability in key function-related loci may help dCGF learn the species genotype-community function landscape. This could be achieved by rational selection of the training communities or by using high-throughput genetic perturbations techniques to generate data, such as CRISPR [70] and genome-wide transposon libraries [71].

Sensitivity analysis of dCGF provided some mechanistic insights into butyrate fermentation that are consistent with the literature. As with any explainable AI method, these results must be interpreted carefully, as the model may have learned a statistical relationship between GFs and function, instead of fully revealing the intra-cellular regulations and inter-species interactions. For instance, *Anaerostipes caccae* (AC) is the highest butyrate producer in the butyrate dataset across a broad range of communities [27]. AC harbors unique GFs, including formaldehyde oxidation and assimilation pathways, that are not present in any other training species. Therefore, the model learns that these features contribute substantially to butyrate production (**Figure 4**B). However, AC’s strong butyrate fermentation capability may arise from complex interactions among GFs that the model did not manage to learn, or from un-annotated features that are not present in its GF vector. To further determine the impacts of these unique GFs in AC on community function, communities including closely related species lacking these features or communities containing mutants of these loci in AC could be characterized and the data used to inform dCGF. Moreover, the collinearity between community functions, such as the negative correlation between butyrate and lactate, creates challenges to identify GFs uniquely contributing to a function.

The encoding of features is a critical determinant of the insights that can be generated and the prediction performance of dCGF. For all except the nitrate and nitrite datasets, we directly used the presence/absence of metabolic pathways from BioCyc as the GFs. While some expert knowledge from the literature may be leveraged to narrow down the most relevant metabolic pathways for each dataset to improve model performance, we did not perform such feature selection. Therefore, the model performances we reported reflect dCGF in the “default” setting. Moreover, for many species, BioCyc annotations are generated automatically using bioinformatic tools and have not been manually curated [72], which may introduce mis-annotated or un-annotated GFs. Finally, additional realizations of the dCGF modeling framework are possible with alternative ML model implementations. For example, the encoders and decoders of dCGF may be implemented with random forest models instead of neural networks. In addition, depending on data availability, set transformers [73], which is encompassed in the broader class of deep set ML models, may capture interactions among GF more effectively to improve prediction performance.

Overall, we present dCGF as a proof-of-concept data-driven modeling framework to learn species genotype to community function mapping in synthetic microbial communities. We demonstrated that dCGF can accurately predict functions of new communities assembled either partially or entirely from new species with known GFs absent from training data. These results show that dCGF is a promising new avenue to explore the immense community design space and to learn useful genotype-function mapping information from data. This new capability can be leveraged to address pressing societal needs in health, agriculture, and environment via accelerating the design-test-learn cycle of synthetic microbial communities with tailored and optimized functions.

## Acknowledgements

We would like to thank Anthony Gitter, Philip Romero, Jaron Thompson, and Alfred Hero for helpful suggestions and feedbacks. This work was funded by Multi-University Research Initiative from the Army Research Office grant W911NF-19-1-0269 (Y.Q. and O.S.V.), the National Institutes of Biomedical Imaging and Bioengineering grant under grant number R01EB030340 (Y.Q. and O.S.V.), R35GM124774 (Y.Q. and O.S.V.), the Defense Advanced Research Projects Agency grant HR0011-23-2-0001 (Y.Q. and O.S.V.), the National Science Foundation under grant number 2448203 (Y.Q., S.D.M. and O.S.V.)and the National Institute of Diabetes and Digestive and Kidney Diseases of the National Institutes of Health under award R01DK133468 (N.Q.-B and S.M.G.).

## Author contributions

Y.Q. and O.S.V. conceived of the study, designed the figures, and wrote the manuscript. Y.Q. developed the methodology, wrote custom scripts to implement the dCGF model, and curated published experimental data. Y.Q. and S. D. M. performed analysis using dCGF model. N.Q.-B. performed GEMs-FBA simulations and S.M.G. interpreted the simulation results. O.S.V supervised the study and O.S.V. and S.M.G. secured funding.

## Data availability

Customized PyTorch modules for dCGF and an example Jupyter Notebook containing basic examples to train and test dCGF are available as a Supplementary File to this manuscript. The code will be made available from https://github.com/VenturelliLab at the time of publication.

## Methods

### Model implementation, hyperparameter selection, and training

The dCGF models were implemented in Python using customized PyTorch [74] modules. Model training was carried out using CPU only on a Mac machine with Apple M1 chip with 16G RAM or on a Mac machine with 2.7GHz Dual-Core Intel i5 chip with 8G RAM. The decoder *y* = *σ*(*z*) in dCGF is a fully connected feed-forward neural network with a single hidden layer of 100 nodes and the encoder for dCGF-IS *z*_*i*_ = *α*(*u*_*i*_, *w*) is a fully connected feed-forward neural network with 200 nodes in the first hidden layer and 30 nodes in the second hidden layer. The community embedding *z* is therefore 30-dimensional. All neural networks use rectified linear unit (ReLU) as activation functions. An Adam solver [75] with learning rate 10^−4^ was applied to optimize the neural network parameters with mean squared error loss. The models were trained for 100 epochs using mini-batch gradient descent. Each mini-batch include no more than 6 communities of identical sizes. Unless stated otherwise, the hyperparemeters above, including the number of nodes in the neural networks, the learning rate, and the training epochs, were used universally on all datasets. Model performance with different hyperparameters can be found in **Supplementary Figure 4** and **Supplementary Figure 5**. No hyperparemeter tuning was performed for specific communities, functions, or evaluation procedures.

### Processing of published data

#### Datasets

Experimental data from human gut microbial communities [27], [46] and soil microbial communities [36], [47] were used to train and test dCGF (**Figure 3**A). Specifically, Clark et al [27] assembled 576 communities *in vitro* from 25 diverse human gut microbial species representing the human gut microbiome. The concentrations of health-related short-chain fatty acids (SCFAs) acetate, butyrate, lactate, and succinate were measured at 48 hrs after innoculation and regarded as community functions. Gowda et al [36] measured the de-nitrification dynamics of converting nitrate 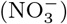 to nitrite 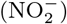 in 186 communities assembled *in vitro* from 12 soil microbes. We used nitrate and nitrite concentrations 8 hr after inoculation in SDM medium as community function. This is a time point when both nitrate and nitrite concentrations are non-zero for most of the communities in the dataset. The Sanchez-Gorostiaga et al [47] dataset measured the rate at which 53 communities assembled from 6 soil microbes degrade starch into simple sugars. Finally, Hromada et al [46] studied the ability of commensal human gut microbial communities assembled from 14 species with different initial species proportions to inhibit human gut pathogen *Clostridioides difficile* (*C. difficile*). Due to the low abundance of *C. difficile* in many samples, the log-transformed relative abundance of *C. difficile* was considered as community function. When multiple replicates were available in the original datasets, we took the average of the replicates.

#### Initial abundances

For the nitrate, nitrite, and starch degradation datasets [36], [47], communities were assembled with equal absolute abundances of all community members. Hence, we assumed unity relative initial abundances *a*_*i*_ = 1 for all species. For the acetate, butyrate, lactate, and succinate datasets [27], communities members were inoculated with equal relative abundances and different communities have identical total abundances. Hence, we assumed initial abundance *a*_*i,j*_ = 1*/n*_*j*_ for each species *i* in community *j*, where *n*_*j*_ is the number of species in community *j*. For the *C. difficile* dataset, experimentally measured initial abundances were used as *a*_*i*_ values.

#### Data standardization

Data standardization is a widely adopted practice to speed up training of machine learning models [76]. It also allows us to compare training performances across different datasets, which have largely different community function outputs in terms of absolute magnitudes, as well as using a same set of training hyper-parameters (e.g., training epochs, learning rates etc.) for all datasets. Community function data in all datasets were standardized by first subtracting the mean and then dividing by the standard deviation before being used for training/testing. That is, before training and testing, all community function data were standardized using the following method:

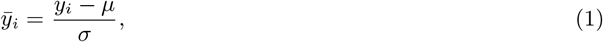

where *µ* = ∑_*i*_ *y*_*i*_*/n*_samples_ and 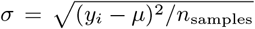 are the mean and the standard deviation of the entire dataset. We considered an alternative standardization scheme, where the mean (*µ*_train_) and the standard deviation (*σ*_train_) of the training data were first computed to standardize the training data using 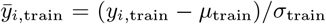. The trained dCGF was then used to predict community function testing data using 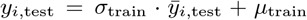, where 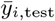 is the dCGF prediction of the testing data. We demonstrated that the *k*-fold cross-validation performances of dCGF were insensitive to data standardization schemes (**Supplementary Figure 13**).

#### Genetic features

The GFs for the species in the acetate, butyrate, lactate, succinate, starch degradation and *C. difficile* datasets from are the presence/absence of metabolic pathways as annotated by the BioCyc genome database collection [50] (version 24.5). The GFs for the species in the denitrification datasets from [36] are the presence/absence of 17 genes identified to be critical to denitrification process in [36]. Strain information and links to each species’ most updated BioCyc databases can be found in Supplementary Table 1. In BioCyc, the presence/absence of metabolic pathways, reactions, genes, and other genetic features in each species is tabulated, which were obtained initially using bioinformatic tools from the species’ reference genomes on NCBI and some species underwent manual curation based on the literature [50]. We did not perform any additional bioinformatic analysis. In BioCyc, each metabolic pathway is assigned a unique ID that is universal in all species. This allows us to compare the presence/absence of metabolic pathways in different species. Metabolic pathway *x* present in species *s*_1_ but absent in *s*_2_, for example, will be included in GF vectors of all species and encoded as 1 (0, respectively) in the GF vector of *s*_1_ (*s*_2_, respectively). For the acetate, butyrate, lactate, and succinate datasets, since the experiments were carried out in defined medium, pathways involving substrates that are not present in the medium were removed from genetic features. These pathways include, for example, xylose degradation, glycogen degradation, chitin degradation, and choline degradation.

### Model performance evaluation

Three procedures were used to evaluate the performance of dCGF in this paper: (i) *k*-fold cross validation (**Figure 3**B), (ii) “leave-one-out” test (Figure **Figure 5**A), and (iii) “‘stringent-leave-out” test (**Figure 5**D). Performances were either evaluated as Pearson correlation between predicted and measured community functions or as root-mean-square error (RMSE), computed as:

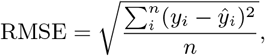

where *y*_*i*_ and *ŷ*_*i*_ are measured and model predicted community functions, respectively, and *n* is the number of samples.

#### *k*-fold cross-validation

To carry out *k*-fold cross validation, a dataset consisting of *n* communities *C*_1_, …, *C*_*n*_ and their measured functions *y*_1_, …, *y*_*n*_ was split randomly into *k* folds, where (*k* − 1) folds of data were used for training and one fold of data was used for testing. This was repeated *k* times until each fold becomes testing data exactly once. The values of *k* were chosen between 6-12 (see **Supplementary Table** 3), depending on dataset size, to allow the folds to be of equal or nearly identical sizes. We also performed analysis using different fraction of the datasets as training data (**Supplementary Figure 2**). To generate **Figure 3**B-C, the above *k*-fold cross validation procedure was repeated 5 times using different random seeds to split the folds and to initialize neural network parameters in dCGF. In panel B, for each community *i* and each random seed *j*, we recorded the predicted community function *ŷ*_*i,j*_ and averaged across the random seeds to make a prediction 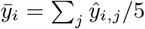. The correlation between 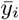 and the measured functions *y*_*i*_ was to generate the scatter plots in **Figure 3**B and to quantify Pearson correlations. In panel C, for each random seed *j*, the correlation between *ŷ*_*i,j*_ and *y*_*i*_, was computed. This was repeated for all 5 random seeds to generate the dots in the bar plot in **Figure 3**C.

#### “Leave-one-out” test

To carry out leave-one-out test using species *i* as testing species, all communities in the dataset that does not include species *i* were used to train a dCGF model. All communities in the dataset that include species *i* were left out as testing data and used to evaluate the model’s prediction performance, summarized by Pearson correlation coefficient between predicted and measured function. This procedure was carried out 5 times using different random seeds to initialize dCGF parameters, and the average Pearson correlation coefficient was used to summarize model performance when species *i* is the testing species.

#### “Stringent-leave-out” test

To carry out stringent leave-out test for a dataset including a set of 𝒮 species, communities containing a set of ℒ ⊂ 𝒮 testing species were excluded from training, and only communities including the rest 𝒯 = 𝒮 − ℒ species were used for model training (**Figure 5**D). Data from communities that include species from both sets ℒ and 𝒯 were disposed and data from communities assembled from ℒ were used for testing the performance of dCGF. For the nitrate and nitrite datasets, every possible 4-species combination were used as ℒ once. For the acetate, butyrate, lactate, and succinate datasets, every possible 4-species combinations that leads to more than 10 communities for testing and more than 80 communities for training were used as ℒ once. In both cases, for every (ℒ, 𝒯) partition, stringent leave-out performances were evaluated using dCGF trained from 5 different random initial parameter seeds to generate the distributions in **Figure 5**E and **Supplementary Figure 10**C. The distributions of testing and training sets for different (ℒ, 𝒯) partitions are also shown in **Supplementary Figure 10**A. The other two datasets were not used for stringent leave-out test as they lead to training/testing sample sizes that are too small (e.g., fewer than 10 samples).

#### Comparison with a null model

We compared leave-one-out performance of dCGF with a null model that assumes the testing species has no effect on community function. Hence, suppose community *c*_1_ is composed of a sub-community *c*_0_ and a testing species *x*. Let *y*_*i*_ be the experimentally measured function of community *c*_*i*_, the null model predicts *y*_1_ = *y*_0_. This null model prediction is only available when subcommunity *c*_0_ and community *c*_1_ are both included in the dataset. On the other hand, dCGF’s prediction does not rely on the availability of *y*_0_ in the dataset. In general, given a testing species *x*, let *C*_*x*_ be all the communities in the dataset where *x* is a member species, the communities that the null model can make prediction is a subset of *C*_*x*_, which we denote by *C*_*x*,null_ ⊆ *C*_*x*_. For a given community function with testing species *x*, for dCGF, we used communities in *C*_*x*,null_ to compute the Pearson correlations in **Figure 5**C. The *C. difficile* dataset was excluded from both figures as it is composed of communities with different initial abundances, and thus the null model predictions are not unique.

### Sensitivity analysis

We performed sensitivity analysis on trained dCGF models to identify genetic features that are potentially impactful for a community function. In particular, sensitivity analysis was performed by numerically silencing/turning on a genetic feature in a species. First, all available experimental data were used to train an ensemble of *n*_*r*_ dCGF models with different random initial parameters: *y* = *F*_*i*_(*u*), *i* = 1, …, *n*_*r*_, where *u* is the genetic feature matrix, *y* is the phenptype, and *F*_*i*_ is a dCGF trained from all available data with the *i*-th initial random seed. The sensitivity of community function *y* to genetic feature *i* in species *j* in model *k* is:

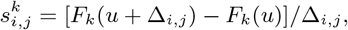

where Δ_*i,j*_ is a perturbation matrix with the same size as *u*. Let *v*_*j*_ represent the initial abundance of species *j*, if genetic feature *i* is present in species *j*, then the (*i, j*)-th element in Δ_*i,j*_ is −*v*_*j*_ and all other elements in Δ_*i,j*_ is zero. This silences genetic feature *i* in species *j*. Conversely, if genetic feature *i* is absent in species *j*, then the (*i, j*)-th element in Δ_*i,j*_ is *v*_*j*_ and all other elements in Δ_*i,j*_ is zero. This turns on genetic feature *i* in species *j*. Sensitivity of the function to genetic feature *i* was then computed as the average from the ensemble of models and all random seeds: 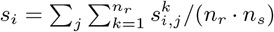, where *n*_*s*_ is the number of species in a community.

### Model pre-training

We used model pre-training to incorporate mechanistic knowledge of short-chain fatty acid biosynthesis, especially those related to butyrate fermentation, into dCGF. The last step of acetyl-CoA to butyrate fermentation involves converting butyryl-CoA to butyrate. This transformation can occur via two pathways: via butyrate kinase and phosphotransbutyrylase (butyrate kinase pathway) or via butyryl-CoA:acetate CoA-transferase (CoA transferase pathway) [52], [77]. The CoA transferase pathway is present in most butyrate producers and is much more efficient than the butyrate kinase pathway [77], [78]. For example, among all species in the butyrate dataset, only *Coprococcus comes* (CC) harbors and uses the butyrate kinase pathway. Additionally, in the CoA transferase pathway, the CoA moiety of butyryl-CoA is transferred to external acetate, forming acetyl-CoA and reducing acetate concentration. Upstream of these reactions, acetyl-CoA is converted from pyruvate via the pytuvate dehydrogenase as the main substrate for butyrate biosynthesis. Some bacteria, such as *Anaerostipes caccae* (AC) and *Desulfovibrio piger* (DP), has the ability to convert lactate into pyruvate [27], [79]–[81]. In AC, this increases the flux of acetyl-CoA to butyrate fermentation. Therefore, these pathways have substantial impacts on the concentration of acetate, butyrate, and lactate.

The butyrate kinase pathway and lactate to pyruvate fermentation pathways are not included in any species in the SCFA datasets by BioCyc. Thus, we first appended them to the genetic feature vectors and then generated 5,000 synthetic species by assigning a random binary number to each genetic feature in each synthetic species. Depending on the presence/absence of the three driver pathways: CoA transferase (pwy 1), butyrate kinase (pwy 2), and lactate to pyruvate fermentation (pwy 3), monoculture of each of the synthetic species is assigned random, correlated values of butyrate, acetate, and lactate concentrations by sampling from uniform distributions on different intervals as synthetic functions (**Supplementary Figure 10**B, **Supplementary Table** 4). Using these synthetic monoculture data, we expect the pre-trained model to learn the qualitative effects of the three driver pathways on SCFAs, while assuming nothing on the impacts of other genetic features on intracellular regulations, inter-species interactions, and SCFA production. These synthetic data were used to pre-train dCGFs for 10 epochs. The pre-training process described above were repeated five times using different random seeds to generate synthetic species’ genetic features, to generate synthetic data, and to initialize NN parameters, leading to an ensemble of 5 different pre-trained dCGFs. Their NN parameters were used as initial parameters for dCGFs when trained on experimental data for 100 epochs. The distributions in **Supplementary Figure 10**C arise from different random seeds as well as different ways to partition training and testing species.

### Synthetic data generation

#### Strategy overview

To demonstrate that dCGF can capture a variety of complexities shaping microbial community functions, which often take the forms of inter-species interactions and intra-cellular interactions, we trained and tested dCGF using synthetic data, which were simulated using ground-truth mechanistic models. We assume that microbial community assembly can be modeled as microbe-effector ODE models, taking the general form:

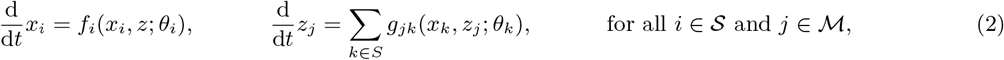

where 𝒮 is the set of all species, ℳ is the set of all molecular effectors, *x*_*i*_ is the abundance of species *i, z*_*j*_ is the concentration of molecular effector *j*, and *θ*_*i*_ is a list of parameters related to species *i*. These parameters quantify, for example, the rate constant that a species consumes a resource and convert it to biomass. The molecular effectors with concentration vector *z* may represent cross-feeding metabolites, resources, or toxins (**Figure 2**F). For instance, a prime example of (2) is the consumer resource model [36], [82]. Assuming *z*_0_ is the only resource, the functions *f*_*i*_ and *g*_0*k*_ take the form: *f*_*i*_(*x*_*i*_, *z*_0_; *θ*_*i*_) = *a*_*i*_*b*_*i*_*x*_*i*_*z*_0_ and *g*_0*k*_(*x*_*k*_, *z*_0_; *θ*_*k*_) = −*b*_*k*_*x*_*k*_*z*_0_, where *a*_*i*_ is the biomass conversion rate constant and *b*_*i*_ is the resource consumption rate constant for species *i*. The function of the community *y* is a dependent on the abundance of all microbes and the concentration of the molecular effectors: *y* = *h*(*x, z*). To map the genotype of a species *i* to community function, we assume that the microbe-effector model parameters associated with species *i* are deterministic functions of its genotype *v*_*i*_: *θ*_*i*_ = (*v*_*i*_). As a consequence, the solution to the microbe-effector model, *x*(*t*) and *z*(*t*), and consequently the community function *y* = *h*(*x, z*), are functions of the genotype. This modeling strategy to map genotypes to functions have been applied in [36], where (2) is a consumer-resource model and ϕ (·) is a linear function.

#### Microbe-effector ODE model

To generate the synthetic data in **Figure 2**G and **Supplementary Figure 1**, the microbe-effector ODE model describes the dynamics of species’ absolute abundances *x*_*i*_ as well as inter-species interactions mediated through molecular effectors: a polysaccharide *z*_1_, a polysaccharide break-down product *z*_2_, a monosaccharide *z*_3_, a cross-feeding growth factor *z*_4_, two toxins *z*_5_ and *z*_6_, and a metabolite output *y* that contributes to community function. The species growth dynamics generally follow:

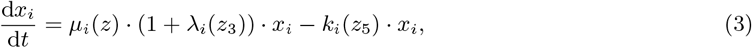

where *µ*_*i*_(*z*_1_, *z*_2_, *z*_3_) is the growth rate of species *i* that depends on the concentrations of the molecular resources *z*_1_, …, *z*_3_, *λ*_*i*_(*z*_4_) is the fraction increase in growth rate due to the presence of the cross-feeding growth factor (*z*_4_), and *k*_*i*_(*z*_5_) is the killing rate constant of species *i* due to apoptosis and the presence of the toxin *z*_5_. Specifically, we define:

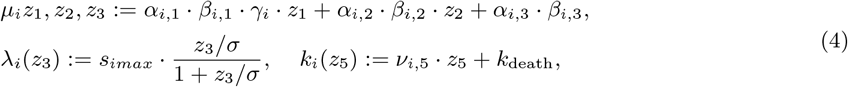

where α_*i,j*_ describes the rate species *i* consumes resource *j*, β_*i,j*_ describes the rate species *i* converts resource *j* into biomass, *s*_*max*_ is the maximum amount of growth rate stimulation provided by cross-feeding metabolite *z*_3_, *σ* is the EC50 of *z*_3_, ν_*i,j*_ is the killing rate constant of toxin *z*_*j*_, and *k*_death_ is a fixed parameter describing species’ apoptosis rate. For species *i* that can degrade and utilize the polysacchride *z*_1_, a fraction γ_*i*_ of it is consumed by *i* for its growth and the rest (1 − *γ*_*i*_) fraction of the resources are degraded, typically by outer-membrane fiber degradation enzymes [83], and then released into the environment. The polysacchride *z*_1_ and monosaccharide *z*_3_ can be consumed by community members. Hence, their dynamics follow:

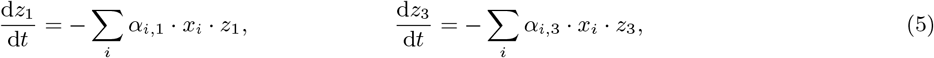

The cross-feeding growth factor *z*_4_, toxins *z*_5_ and *z*_6_, and output metabolite *y* are produced by species *i* with constant rates *η*_*i*,4_, *η*_*i*,5_, and φ_*i*_ respectively:

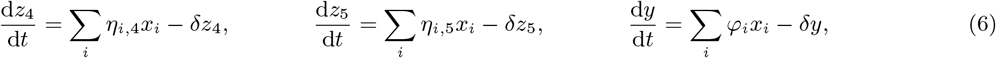

where *δ* is a fixed degradation rate constant for the molecular effectors. The polysacchride break-down product *z*_2_ is both produced and consumed by community members:

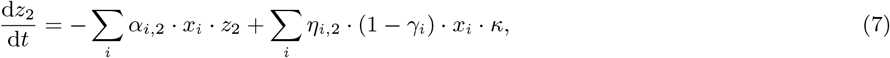

where κ is a fixed parameter describing the molar conversion rate between the polysacchride and the its break-down product, as a polysacchride molecule can be degraded into multiple monosacchride molecules. Equations (3)-(7) are applied to every species and molecular effectors in the community. For species that cannot produce (or consume) a molecular effector, the related parameters are set to 0. For example, if species *i* is unable to consume resource *j*, the consumption rate constant α*i, j* is set to zero. Similarly, if it is unable to produce toxin *z*_5_, the production rate constant *η*_*i*,5_ is set to zero. We determine these rate constants based on the GF vectors of each species using the rules outlined below.

#### Model parameter generation

To determine the parameters related to each species, we assume that each parameter is impacted by a small number of GFs in the species’ GF vectors. Specifically, for every parameter θ_*i*_ ∈ {*α*_*i,j*_, *β*_*i,j*_, *η*_*i,j*_, *φ*_*i*_, *γ*_*i*_} related to species *i*, we use the following model to describe the impact of GFs on species’ dynamic parameters (i.e., intra-cellular regulation):

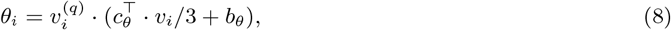

where 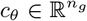 contains three non-zero elements sampled randomly from a non-negative uniform distribution, describing the weight of the GFs to parameter *θ*_*i*_, *v*_*i*_ is the *n*_*g*_-dimensional binary GF vector of species *i*, representing the presence/absence of each GF in species *i*, and *b*_*θ*_ is a positive bias. The *q*-th element of the GF vector for species *i*, 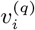 determines if *θ*_*i*_ will be zero or non-zero: if 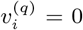, then the parameter *θ*_*i*_ will be zero regardless of the weight vector *c*_*θ*_ and the bias *b*_*θ*_. On the other hand, if 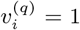, then the magnitude of the parameter *θ*_*i*_ is determined by the three GFs where *c*_*θ*_ has non-zero elements. Note that both *c*_*θ*_ and *b*_*θ*_ are not functions of *i*. This reflects the assumption that there is a universal rule from species’ GF to their interactions with molecular effectors. For example, if a set of four genes determine the rate at which a species x can degrade polysacchride *z*_1_, the same set of genes determine species y’s capability to degrade polysacchride *z*_1_.

#### Model simulation

We generated 40-dimensional binary GF vectors by sampling from the uniform binary distribution for 1010 synthetic species. The first 1000 species were labeled as training species to assemble training communities for synthetic training data generation. The last 10 species were labeled as testing species to assemble testing communities for synthetic testing data generation. This procedure creates a synthetic microbial community dataset for stringent-leave-out test. Specifically, for the training data, we included all mono-species and randomly selected multi-species sub-communities with up to 10 members. To avoid the training data being dominated by large communities, communities of any given size can have no more than 1000 sample. For the testing data, we included all possible sub-communities that can be assembled from the 10 testing species. Based on the microbe-effector model and the parameter assignment rule described above, we simulated both the training and testing communities till time=48 and the output metabolite concentration *y* is used to quantify community function. The procedure described above was applied 10 times using different random seeds to generate random binary GF vectors *v*_*i*_, to generate random weight vectors *c*_*θ*_, to randomly pick sub-communities as training communities, and to initialize dCGF parameters. For each random seed used to generate synthetic community function data, we trained the model using 5 different random seeds to initialize parameters in the neural networks of dCGF-IS and dCGF-ES.

### Species functional embedding

With reference to **Figure 6**A, to derive the functional embedding of species *i* for a community function *y*, we first trained dCGF using all available data for community function *y*. The encoder part of the trained dCGF was saved, and we denote it as a mapping *E*(·). In order to compare species’ functional embeddings without the impact of their abundance, we assumed all species have unity abundance when computing their functional embeddings. Hence, we compute each species’ functional embedding using *z*_*i*_ = *E*(*v*_*i*_), where *v*_*i*_ is the binary GF vector for species *i*. To compare species’ functional similarities using their genetic features, we performed principal component analysis (PCA) on the matrix 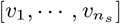 with dimension *n*_*g*_ × *n*_*s*_, where *n*_*g*_ is the number of GFs and *n*_*s*_ is the number of all species appearing in a dataset. To compare species’ functional similarities using their functional embeddings, we performed PCA on matrices 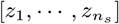 with dimension *n*_*e*_ × *n*_*s*_, where *n*_*e*_ is the dimension of the species’ functional embeddings. To derive species’ functional embeddings using random genetic features (**Supplementary Figure 12**), the binary GF vector *v*_*i*_ for each species *i* was replaced by a random binary vector drawn from the discrete uniform distribution and the steps described above were repeated. PCA was performed using PCA module from the scikit-learn library.

### GEMs-FBA model simulation using MICOM

To perform GEMs-FBA simulations for the acetate, butyrate, lactate, and succinate datasets, we used MICOM software package [42]. Taxonomic abundance data from 16S amplicon sequencing was aggregated at the genus level for all samples. Microbial community metabolic models (MCMMs) were constructed for each sample using MICOM v.0.33.0 [42] with the build() function, excluding taxa below 0.1% relative abundance. Genus-level GEMs from the AGORA database (v.1.03) provided taxonomic reconstructions for these models. Using the defined medium (DM38) used for *in vitro* experiments as a scaffold, each component was matched to the Virtual Metabolic Human database (www.vmh.life). This created an *in silico* medium with flux limits proportional to concentrations, with iron(III) added to achieve a minimum community growth rate of 0.3 hr^−1^ as determined by using `MICOM’s fix medium()` function. The resulting medium is available at https://github.com/Gibbons-Lab/scfa_predictions/tree/main/media. After applying the *in silico* medium to constrain import fluxes, growth rates and metabolic fluxes were calculated using MICOM’s row() function, which employs a cooperative tradeoff flux balance analysis (ctFBA) approach. This method first optimizes community-wide biomass production, then distributes taxon-specific growth rates to maximize individual taxa growth while maintaining a community growth rate close to the optimal. A tradeoff parameter of 0.7, chosen to allow growth for over 90% of taxa, was applied to balance individual and community growth. All models achieved the minimum community growth threshold of 0.3 hr^−1^. Finally, metabolic production data from each MCMM was gathered using the `production rates()` function.

