## Supplementary Information for "A data-driven modeling framework for mapping genotypes to synthetic microbial community functions"

### Supplementary Note 1 dCGF-ES is strictly less expressive than dCGF-IS

We define

$$\mathcal{F}_{\text{IS}} = \left\{ F_{\text{IS}}(\cdot) : \mathbb{R}^{n_g \times n_s} \mapsto \mathbb{R}^{n_y} \,\middle|\, F_{\text{IS}}(u) = \sigma \left[ \sum_i \alpha(u_i, w) \right] \right\},$$

$$\mathcal{F}_{\text{ES}} = \left\{ F_{\text{ES}}(\cdot) : \mathbb{R}^{n_g \times n_s} \mapsto \mathbb{R}^{n_y} \,\middle|\, F_{\text{ES}}(u) = \sigma \circ \alpha \left( \sum_i u_i \right) \right\}$$

as the sets of functions mapping from GF matrix $u$ to community function $y$ arising from dCGF-IS and dCGF-ES, respectively, where $\sigma$ and $\alpha$ are arbitrary real functions with appropriate dimensions, $u_i$ is the $i$-th column of GF matrix $u$, and $w = \sum_i u_i$. We provide a proof sketch to show that the set $\mathcal{F}_{\text{IS}}$ is strictly larger than $\mathcal{F}_{\text{ES}}$.

**Claim**: $\mathcal{F}_{\text{ES}} \subsetneq \mathcal{F}_{\text{IS}}$.
**Proof**: This claim can be shown in two steps.

1. dCGF-ES is no more expressive than dCGF-IS: $\mathcal{F}_{\text{ES}} \subseteq \mathcal{F}_{\text{IS}}$.
   - This part can be shown by construction. In particular, every $f(u) = \sigma \circ \alpha(\sum_i u_i) \in \mathcal{F}_{\text{ES}}$ satisfies $f \in \mathcal{F}_{\text{IS}}$, since it can be written as $f(u) = \hat{\sigma}(\sum_i \hat{\alpha}(u_i, w))$, where $\hat{\sigma}(x) = \sigma \circ \alpha(x)$ and $\hat{\alpha}(x, y) = x$.
2. There exists $f^* \in \mathcal{F}_{\text{IS}}$ such that $f^* \notin \mathcal{F}_{\text{ES}}$.
   - This part can be shown by constructing an example. Let

     $$u_0 = \begin{bmatrix} 1 & 0 \\ 0 & 1 \end{bmatrix} \qquad \text{and} \qquad u_1 = \begin{bmatrix} 1 \\ 1 \end{bmatrix}.$$

     There exists a function $f^* \in \mathcal{F}_{\text{IS}}$ such that $f^*(u_0) \neq f^*(u_1)$. One possible realization is $f^*(u) = \sigma^*(\sum_i \alpha^*(u_i))$

     $$\alpha^*(x, y) = x_1 + x_2 + x_1 \cdot x_2, \qquad \text{and} \qquad \sigma^*(x) = x.$$

     However, for any function $g \in \mathcal{F}_{\text{ES}}$, one necessarily has $g(u_0) = g(u_1)$. Therefore, $f^* \notin \mathcal{F}_{\text{ES}}$.
   - Functions living in the set $\mathcal{F}_{\text{IS}} \setminus \mathcal{F}_{\text{ES}}$ represent community genotype-function mappings that **can** be described by dCGF-IS but **cannot** be described by dCGF-ES. We can develop a better idea of these functions by generalizing the example above.
   - In particular, given two matrices $a, b \in \mathbb{R}^{n,m}$, we denote $a \sim b$ if there exists a column permutation operation pem$(\cdot)$ such that pem$(a) =$ pem$(b)$. Conversely, we denote $a \nsim b$ if for any column permutation operation pem$(\cdot)$, pem$(a) \neq$ pem$(b)$. Suppose that the dimension of the community embedding is sufficiently large, then for any community GF matrices $a \nsim b$ constructed from a finite pool of species, one can always construct a function $f^* \in \mathcal{F}_{\text{IS}}$ such that $f^*(a) \neq f^*(b)$ for every $a, b$. In contrast, for any $g \in \mathcal{F}_{\text{ES}}$, one will have $g(a) = g(b)$ if $a \nsim b$ but $\sum_i a_i = \sum_i b_i$. Hence, $f^* \notin \mathcal{F}_{\text{ES}}$.
   - Connecting mathematical results to biology: a frequent source when $\sum_i a_i = \sum_i b_i$ but $y_a \neq y_b$ is when there is intra-cellular regulations or inter-species interactions.

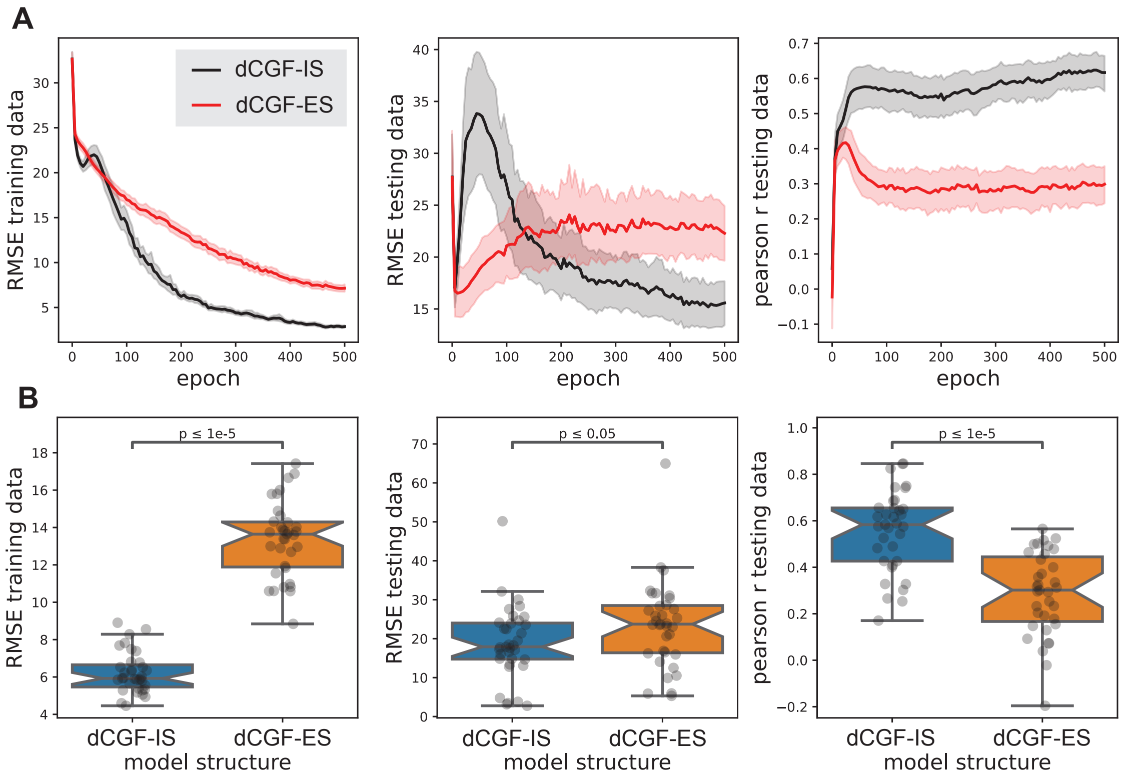


Supplementary Figure 1: **Comparison of dCGF-IS and dCGF-ES performances on synthetic datasets**. (A) Line plots for RMSE of training communities, RMSE of testing communities, and Pearson correlation of testing communities using dCGF-IS and dCGF-ES at different training epochs. Solid lines represent mean and shaded regions represent 95% confidence intervals arising from 10 different random seeds used to generate synthetic community assembly data and, for each set of randomly generated synthetic data, 5 different random seeds used to initialize neural network parameters in dCGF models. (B) Box plots for RMSE of training communities, RMSE of testing communities, and Pearson correlation of testing communities useing dCGF-IS and dCGF-ES trained with 200 epochs. Each dot represents RMSE/Pearson correlation arising from a pair of random seeds used to generated synthetic community assembly data and to initialized neural network parameters. Whiskers indicate 25% and 75% quantiles and notches represent medians. $p$-value are from one-sided Mann-Whitney U tests.

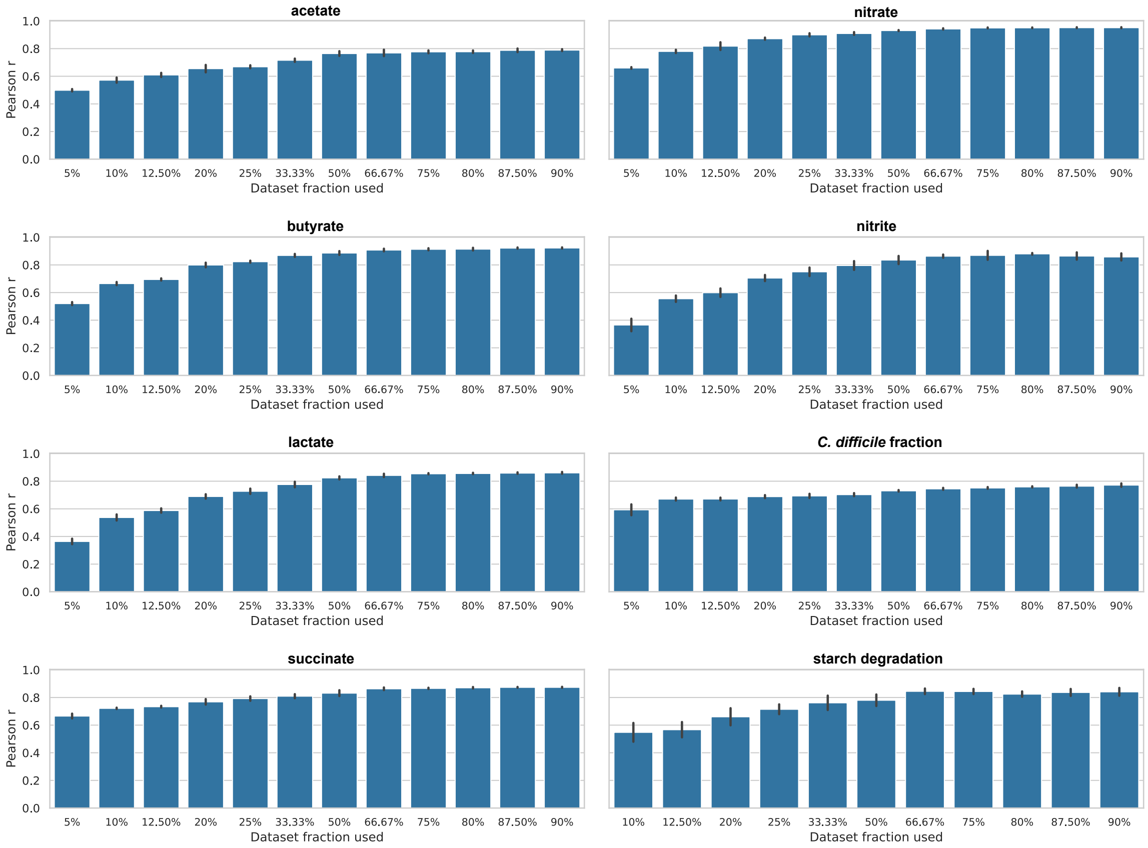


Supplementary Figure 2: **Comparison of $k$-fold cross-validation performances of dCGF-IS using different fractions of data for training**. x-axis labels the amount of experimental data used for dCGF-IS training in each fold. Error bars represent one standard deviations arising from using five random seeds.

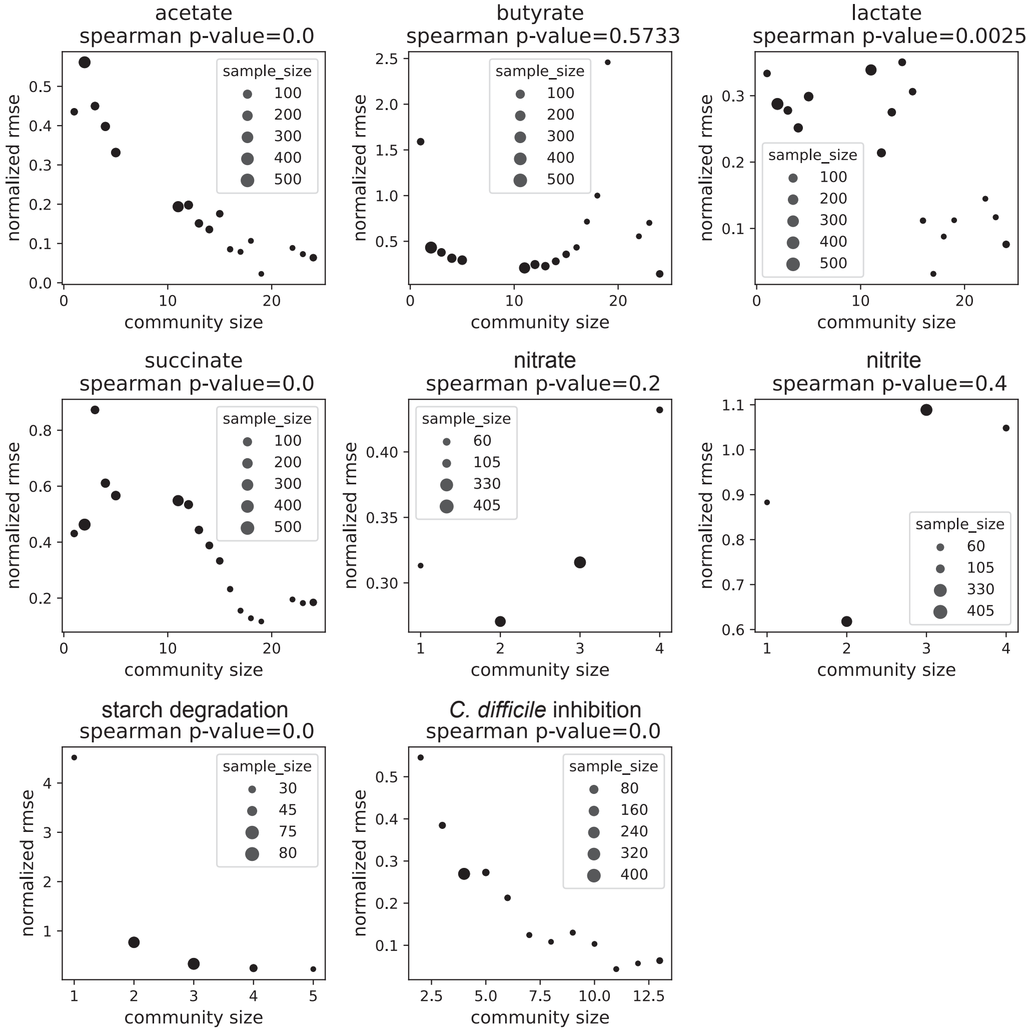


Supplementary Figure 3: **Comparison of $k$-fold cross-validation performances of dCGF-IS for communities of different sizes**. Normalized RMSE for community function $y$ in communities with size $n$ was computed as $\mathrm{RMSE}_{y,n} := \mathrm{RMSE}_{y,n}/\mu_{y,n}$, where $\mu_{y,n}$ is the mean measured function $y$ in all communities with size $n$ and $\mathrm{RMSE}_{y,n}$ is the RMSE error for community function $y$ in all communities with size $n$. To compute RMSE, $k$-fold cross-validation was performed five times for each function using different random seeds to initialize neural network parameters and to split the folds. Each community appears as testing data five times, and the average of the five predictions were used as the prediction of dCGF.

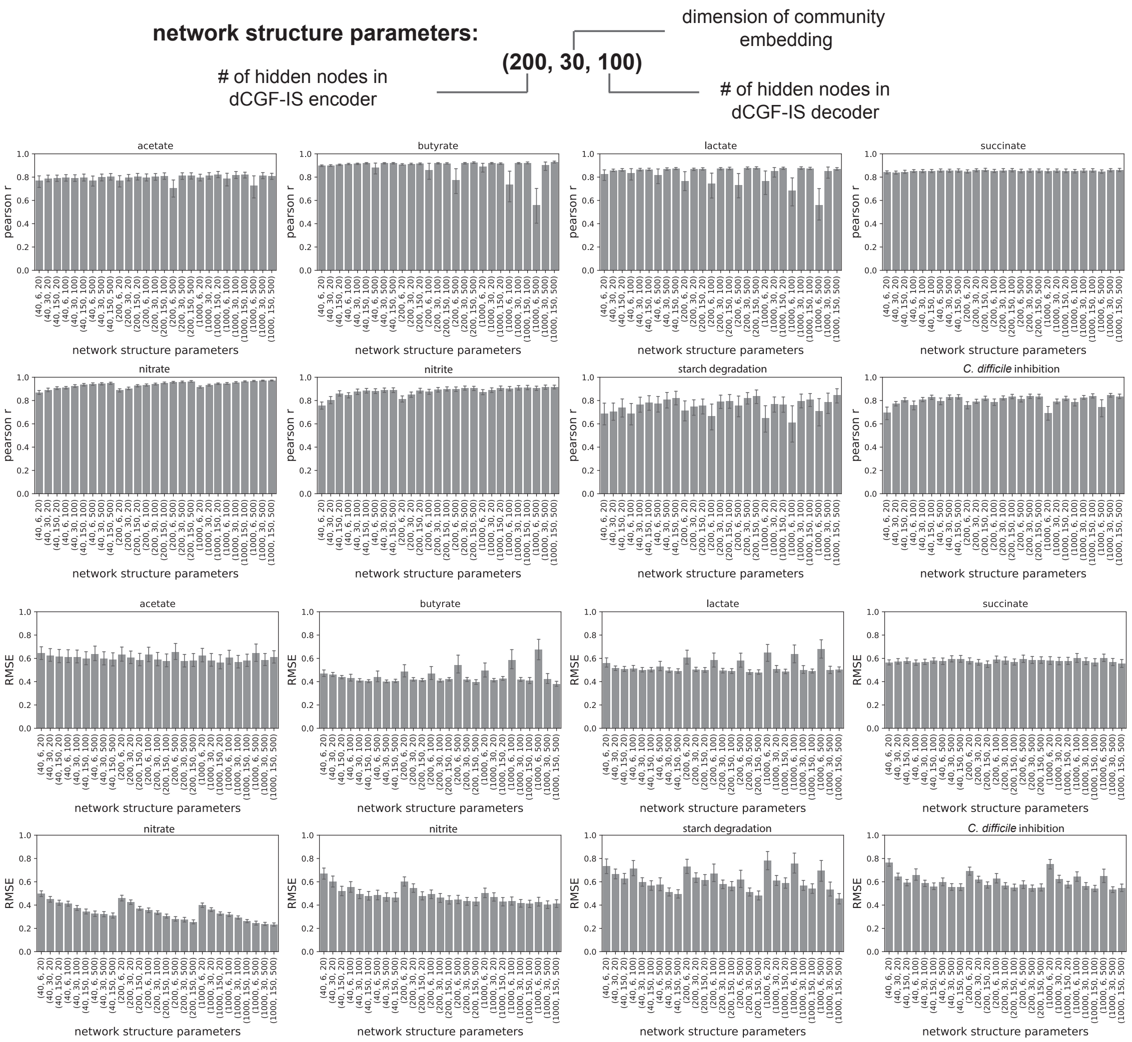


Supplementary Figure 4: **Comparison of $k$-fold cross-validation performances of dCGF-IS using different network structure parameters**. The triplets of numbers on the x-axis represent the number of nodes in the hidden layer of dCGF-IS encoder, the number of outputs from the encoder (i.e., the dimension of the community embedding), and the number of nodes in the hidden layer of dCGF-IS decoder, respectively. Performance were evaluated using (A) Pearson correlation and (B) RMSE. The nominal network structure parameter (200, 30, 100) was used to evaluate model performance in this paper, unless specified otherwise. Performances were evaluated at training epoch=100 using either RMSE or Pearson correlation. Error bars indicate 95% confidence intervals.

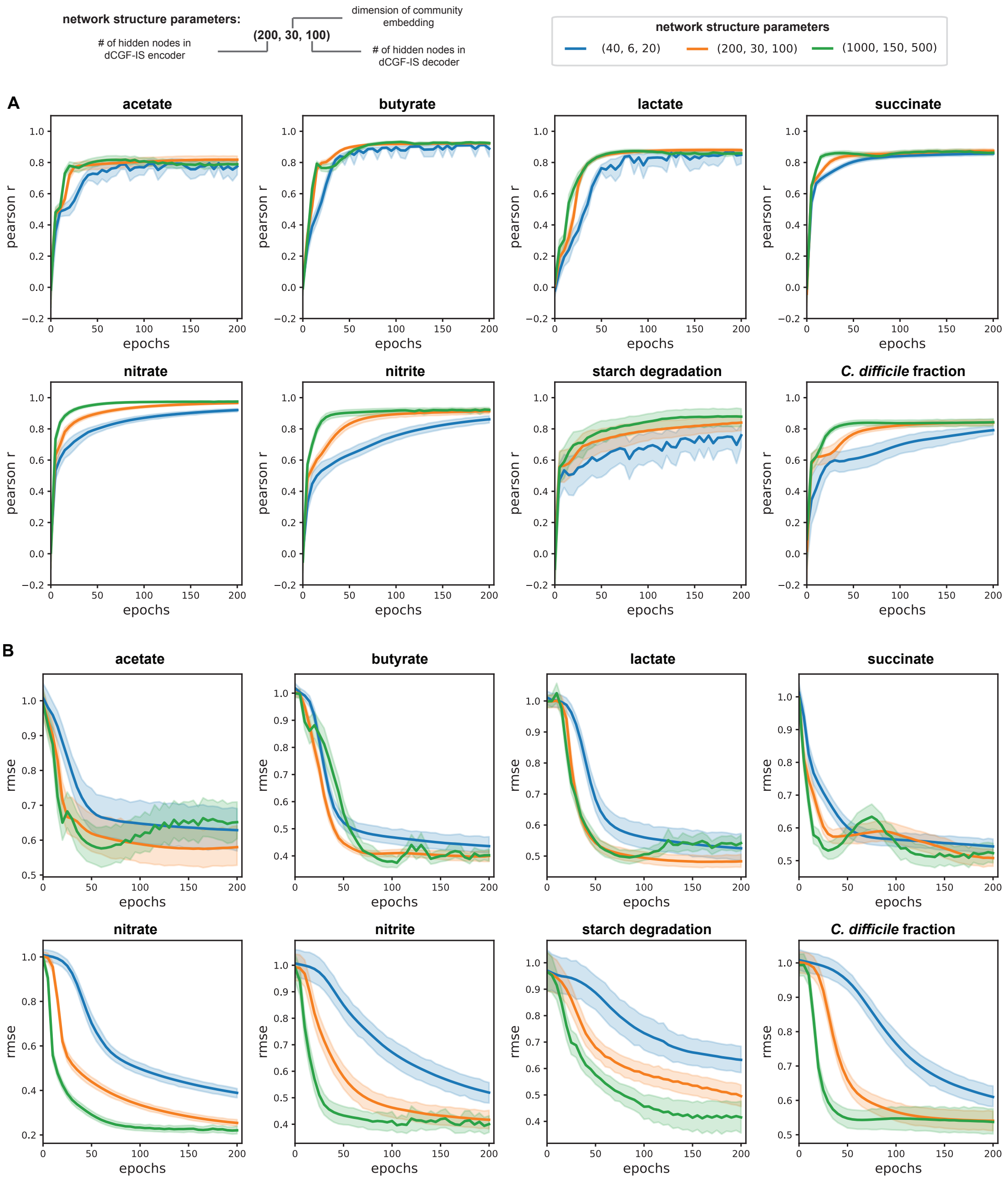


Supplementary Figure 5: **$k$-cross-validation performance of dCGF-IS with different network structure parameters for different training epochs.** Performances were evaluated using (A) Pearson correlation and (B) RMSE. The nominal network structure parameter used to evaluate model performance in the paper is (200, 30, 100), unless otherwise specified. Shaded areas indicate 95% confidence intervals arising from 5 random seeds to split the folds and to initialize neural network parameters.

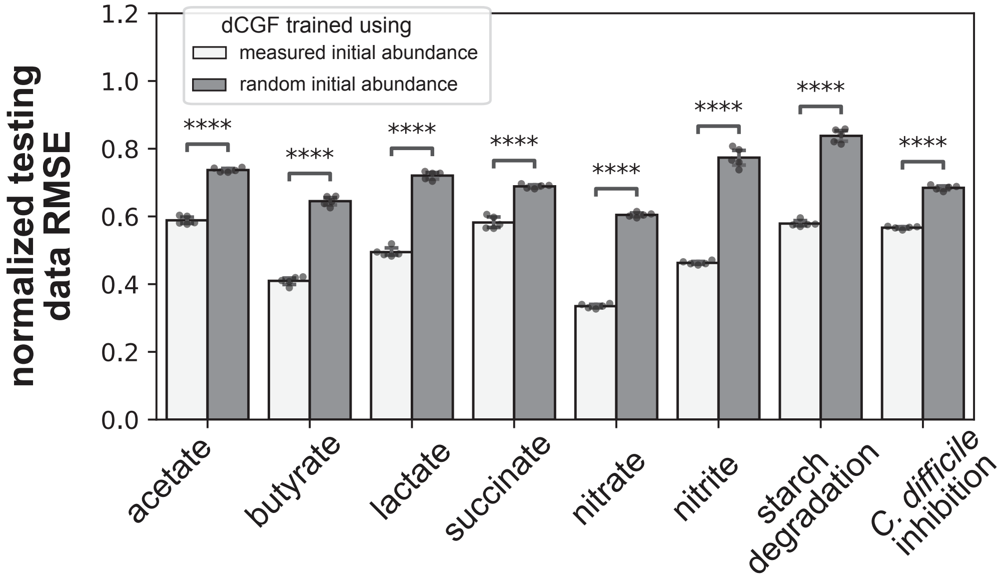


Supplementary Figure 6: **Comparison of $k$-fold cross-validation RMSE of dCGF-IS trained using measured and random initial abundances.** For training and testing using random initial abundances, the initial relative abundances of the community members in each community in the data sets were replaced by a random number drawn from uniform distribution. Each dot in the bar plot represents $k$-fold cross-validation performance using a random initial seed to split the data into folds and to initialize neural network parameters in dCGF-IS. ****: $p < 10^{-4}$ using independent t-test.

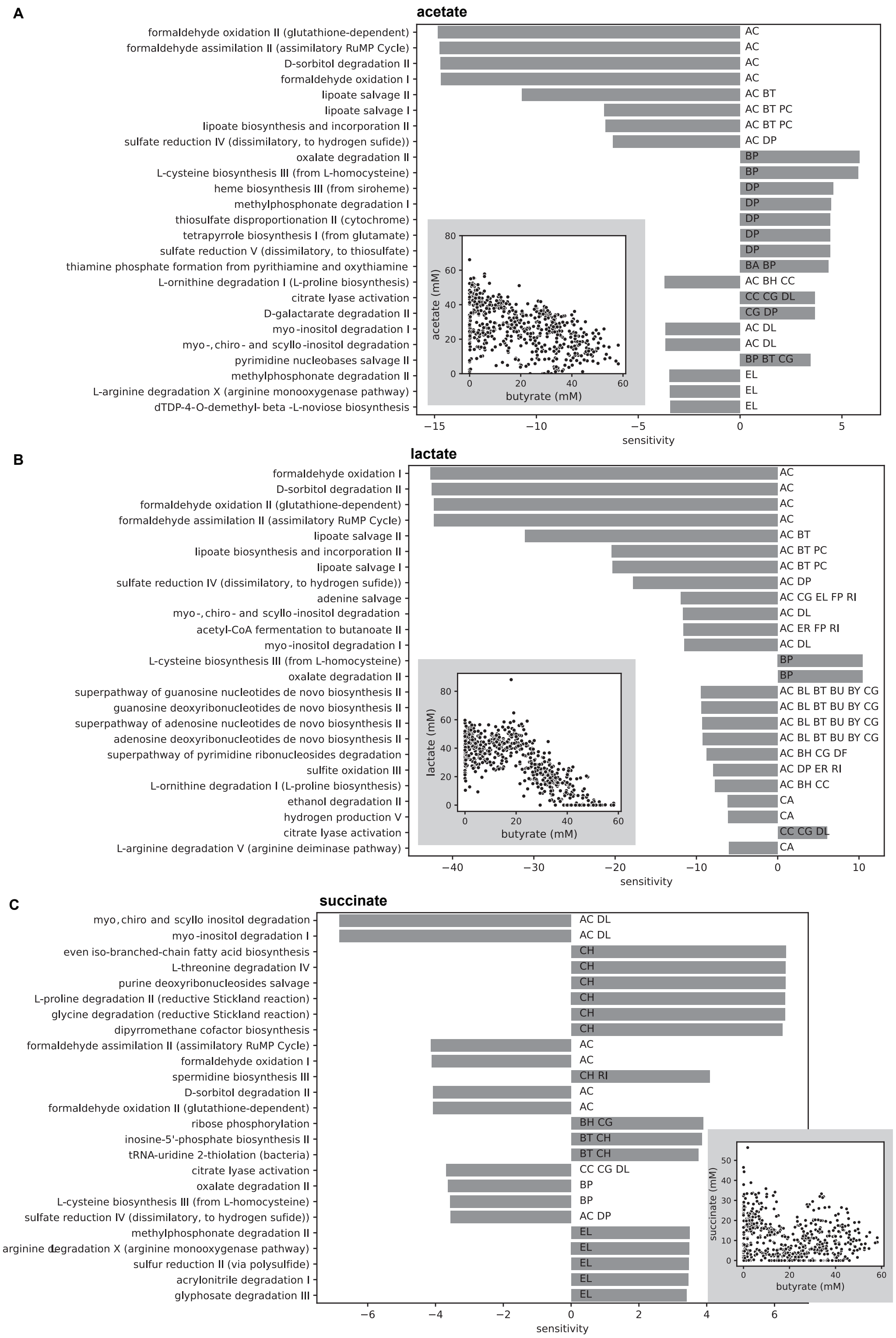


Supplementary Figure 7: **Metabolic pathways with the top 5% sensitivity magnitudes in the acetate (A), lactate (B), and succinate (C) datasets.** Sensitivities were computed for the community composed of all possible species, each with equal initial abundances. Five dCGF-IS models were trained using all data available in the datasets and the sensitivities were computed as the average sensitivities from all five models (see **Methods**). Inset in each panel shows scatter plot of the function with butyrate in experimental data.

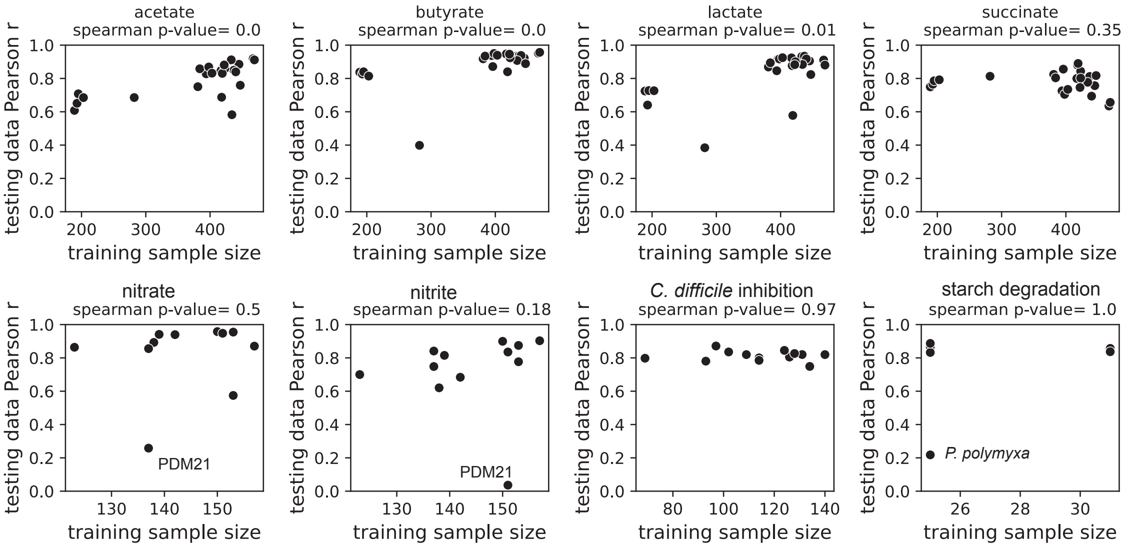


Supplementary Figure 8: **Spearman correlation between leave-one-out prediction performances and training sample size for different testing species**. Each dot represents leave-one-out training data sample size and prediction performance using one species as testing species.

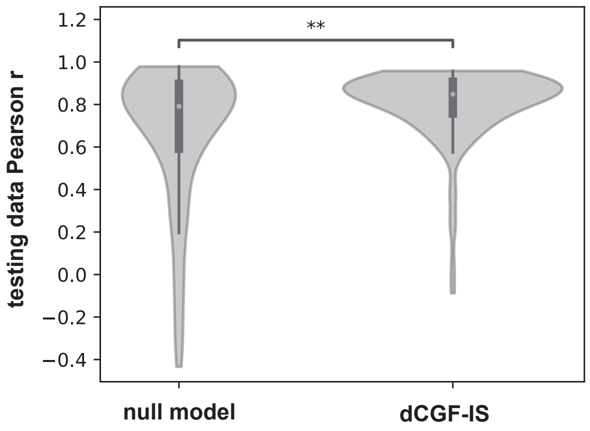


Supplementary Figure 9: **Comparison of leave-one-out prediction performances using dCGF-IS vs. the null model.** Distributions represent the Pearson correlations in leave-one-out tests using different species as testing species (see **Figure 5**C). White dots in the miniature box plots are medians of the distributions. The lower and upper edges of the miniature boxes are the 25% and 75% of the data, respectively. **: $p < 0.01$ using one-sided Mann-Whitney U test.

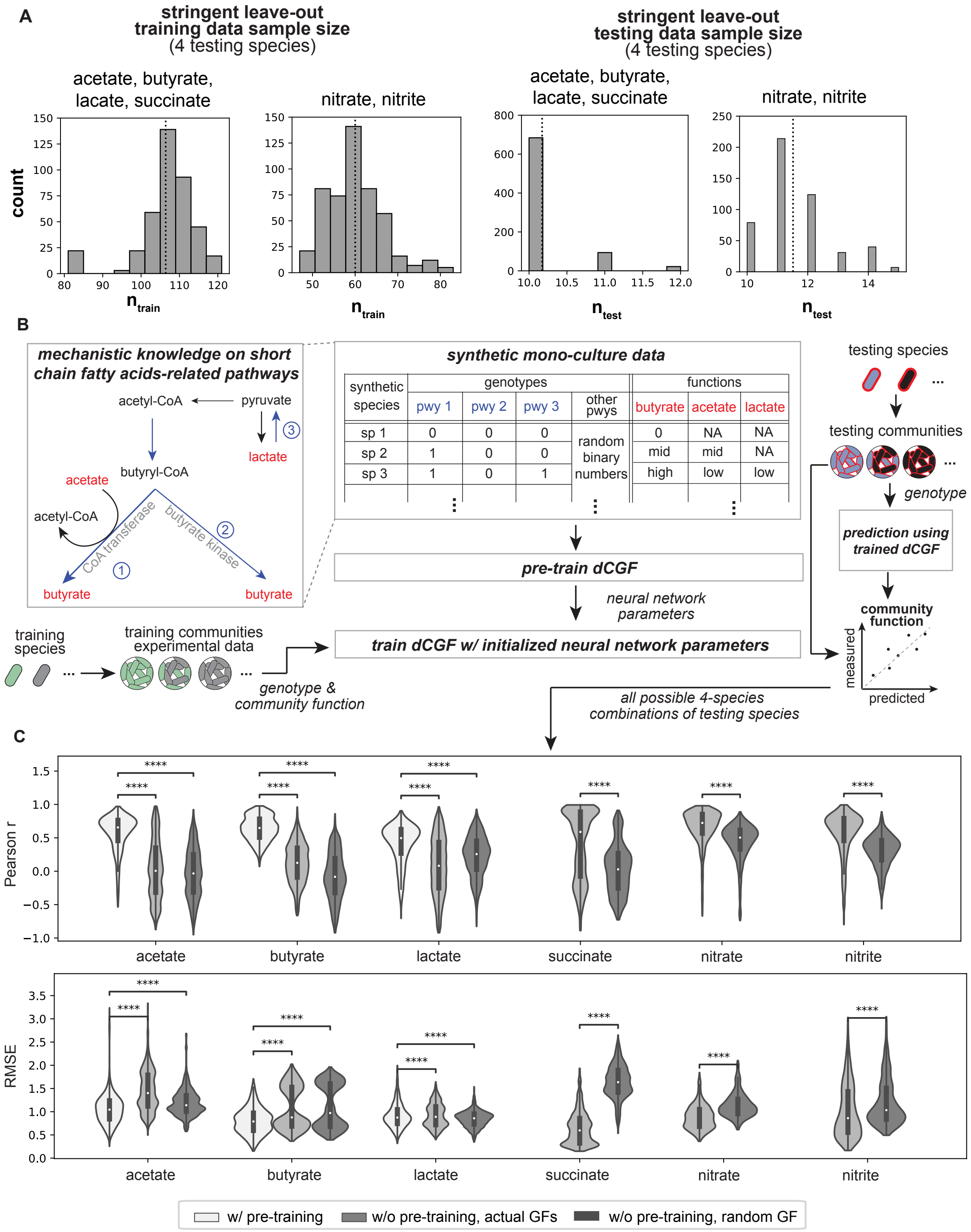


Supplementary Figure 10: Caption on next page

**Pre-training of dCGF-IS improves its stringent-leave-out performance.** (A) Training and testing data sample sizes using 4 species as testing species in stringent leave-one-out tests for different datasets. (B) Illustration of dCGF-IS pre-training procedure. Three key metabolic pathways related to acetate, butyrate, and lactate production were identified. Synthetic mono-culture data were generated by sampling 5000 synthetic species with random GF vectors and assigning acetate, butyrate, and lactate values to these species based on the presence/absence of the three key pathways. These synthetic were then used to pre-train the dCGF-IS model for 10 epochs. The pre-trained model parameters were used to initialize dCGF when given experimental community data for training. See **Methods** for more details. (C) Stringent leave-out performances evaluated by testing community Pearson correlation and RMSE for acetate, butyrate, lactate, succinate, nitrate, and nitrite datasets. Distributions represent performances using four different species as testing species and five different random seeds. The starch degradation and *C. difficile* datasets were excluded for stringent leave-out test because there data sample sizes are too small. Consequently, there does not exist any partition of training/testing species such that there are more than 10 samples for training. ****: $p < 10^{-4}$ using one-sided Mann-Whitney U test.

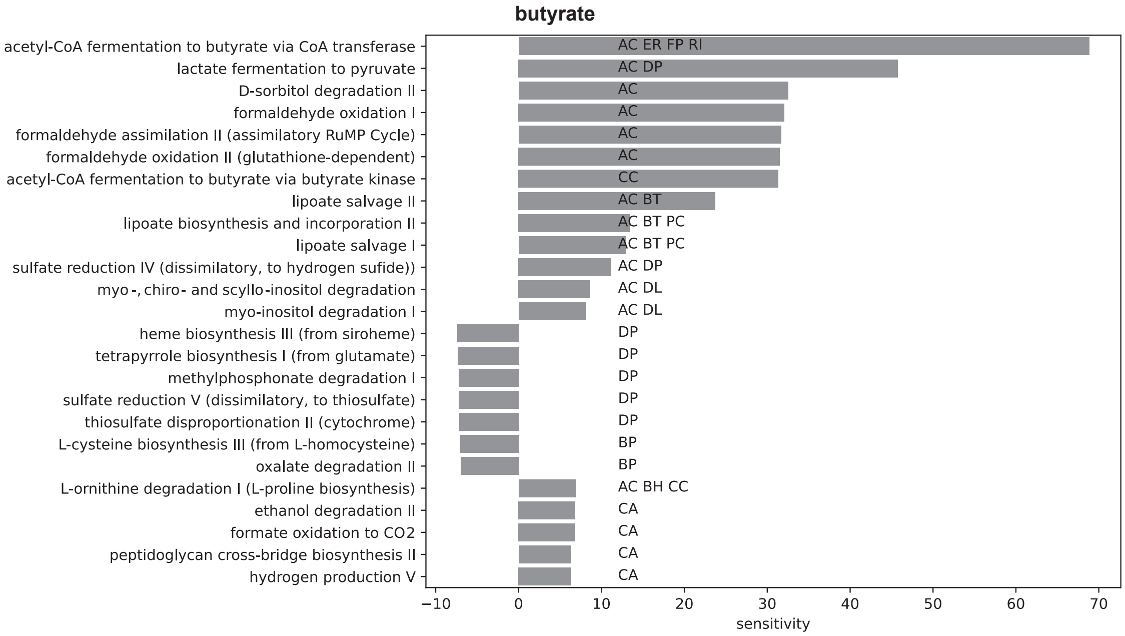


Supplementary Figure 11: **Sensitivity analysis to evaluate the impact of GFs to community butyrate production using pre-trained dCGF model.** All data in the butyrate dataset was used to train the pre-trained dCGF-IS for butyrate. Sensitivity was first computed for each GF in each species and then averaged across all species and all random seeds. See **Methods** for details of model pre-training and sensitivity analysis. Species names on the bars indicate all the species that harbors the corresponding metabolic pathway.

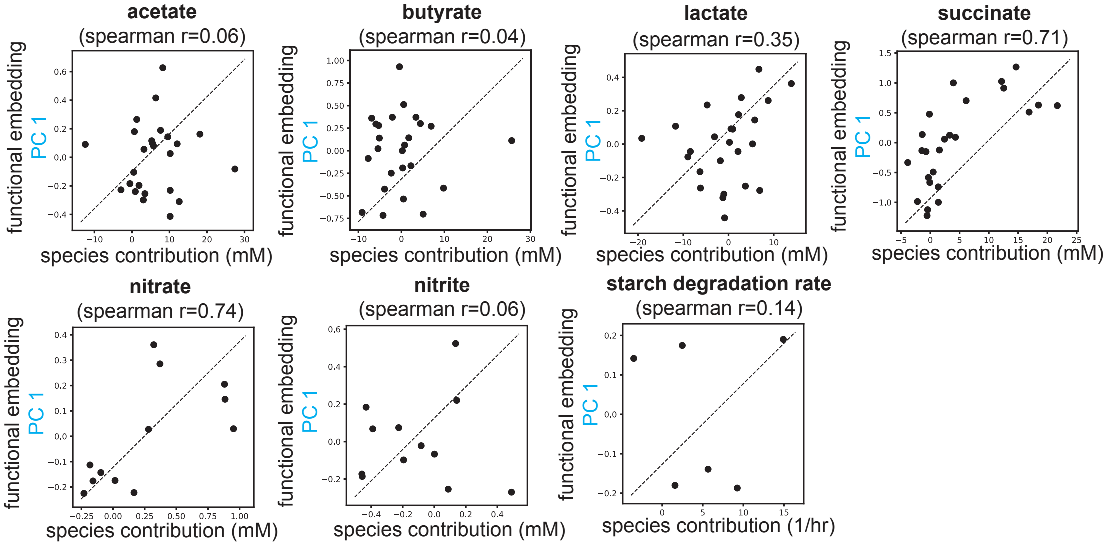


Supplementary Figure 12: **Species' functional embeddings derived from dCGF trained with random GF vectors.**

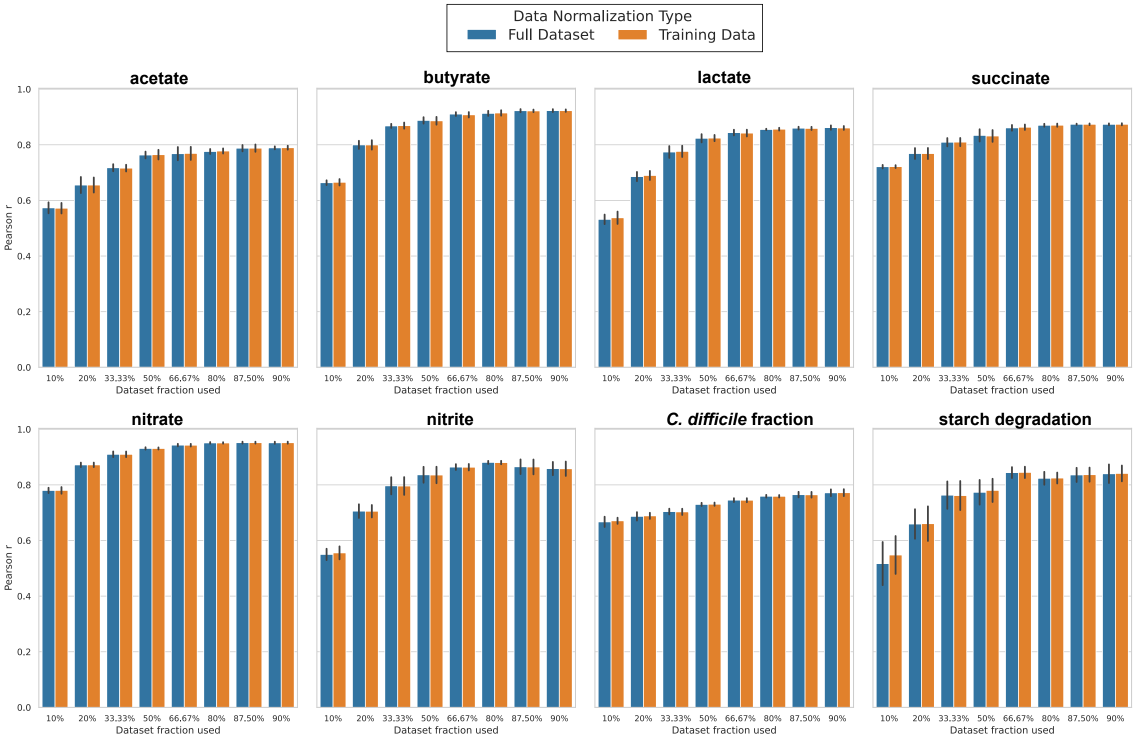


Supplementary Figure 13: **Comparison of $k$-fold cross-validation performances using different data standardization methods.** x-axis labels the amount of experimental data used for dCGF-IS training in each fold for each datasets. Error bars represent one standard deviations arising from using five random seeds. Performances were evaluated as Pearson correlation of the testing data.

| species | strain | abbreviation | reference | biocyc database |
|---|---|---|---|---|
| *Anaerostipes caccae* | DSM 14662 | AC | [27] | link |
| *Bifidobacterium adolescentis* | ATCC 15703 | BA | [27] | link |
| *Bacteroides caccae* | ATCC 43185 | BC | [27] | link |
| *Bacteroides fragilis* | NCTC 9343 | BF | [27] | link |
| *Blautia hydrogenotrophica* | DSM 10507 | BH | [27], [46] | link |
| *Bifidobacterium longum infantis* | ATCC 15697 | BL | [27] | link |
| *Bacteroides ovatus* | ATCC 8483 | BO | [27], [46] | link |
| *Bifidobacterium pseudocatenulatum* | DSM 20438 | BP | [27] | link |
| *Bacteroides thetaiotaomicron* | VPI-5482 | BT | [27], [46] | link |
| *Bacteroides uniformis* | ATCC 8492 | BU | [27], [46] | link |
| *Bacteroides vulgatus* | ATCC 8482 | BV | [27], [46] | link |
| *Bacteroides cellulosilyticus* | DSM 14838 | BY | [27] | link |
| *Collinsella aerofaciens* | ATCC 25986 | CA | [27], [46] | link |
| *Coprococcus comes* | ATCC 27758 | CC | [27] | link |
| *Clostridioides difficile* | R20291 | CD | [46] | link |
| *Clostridium asparagiforme* | DSM 15981 | CG | [27] | link |
| *Clostridium hiranonis* | DSM 13275 | CH | [27], [46] | link |
| *Clostridium scindens* | ATCC 35704 | CS | [46] | link |
| *Dorea formicigenerans* | ATCC 27755 | DF | [27] | link |
| *Dorea longicatena* | DSM 13814 | DL | [27] | link |
| *Desulfovibrio piger* | ATCC 29098 | DP | [27], [46] | link |
| *Eggerthella lenta* | DSM 2243 | EL | [27], [46] | link |
| *Eubacterium rectale* | ATCC 33656 | ER | [27], [46] | link |
| *Faecalibacterium prausnitzii* | A2-165 | FP | [27], [46] | link |
| *Prevotella copri* | DSM 18205 | PC | [27], [46] | link |
| *Parabacteroides johnsonii* | DSM 18315 | PJ | [27] | link |
| *Roseburia intestinalis* | L1-82 | RI | [27] | link |
| *Bacillus cereus* | soil sample | - | [47] | link* |
| *Bacillus megaterium* | ATCC 14581 | - | [47] | link |
| *Bacillus subtilis* | ATCC 23857 | - | [47] | link* |
| *Bacillus thuringiensis* | ATCC 10792 | - | [47] | link |
| *Paenibacillus polymyxa* | ATCC 15970 | - | [47] | link |
| *Paracoccus denitrificans* | ATCC 19367 | PAR19367 | [36] | - |
| *Paracoccus sp.* | PAR01 | PAR01 | [36] | - |
| *Ensifer sp.* | ENS09 | ENS09 | [36] | - |
| *Agrobacterium sp.* | AGB01 | AGB01 | [36] | - |
| *Acidovorax sp.* | ACV02 | ACV02 | [36] | - |
| *Achromobacter sp.* | ACM01 | ACM01 | [36] | - |
| *Psudomonas sp.* | PDM12 | PDM12 | [36] | - |
| *Psudomonas sp.* | PDM13 | PDM13 | [36] | - |
| *Psudomonas sp.* | PDM14 | PDM14 | [36] | - |
| *Psudomonas sp.* | PDM21 | PDM21 | [36] | - |
| *Pantoea sp.* | PNT03 | PNT03 | [36] | - |
| *Psudoxanthomonas sp.* | PXM03 | PXM03 | [36] | - |

Supplementary Table 1: **Microbial strain information.** An asterisk (*) indicates the genome annotation of an alternative strain was used as the original one in the reference was unavailable on BioCyc.

Supplementary Table 2: Leave-one-out performances

| dataset | training species | pearson r | n_train | n_test |
|---|---|---|---|---|
| nitrite | ACM01 | 0.700 | 123 | 63 |
| nitrite | ACV02 | 0.903 | 157 | 29 |
| nitrite | AGB01 | 0.620 | 138 | 48 |
| nitrite | ENS09 | 0.841 | 137 | 49 |
| nitrite | PAR01 | 0.748 | 137 | 49 |
| nitrite | PAR19367 | 0.777 | 153 | 33 |
| nitrite | PDM12 | 0.900 | 150 | 36 |
| nitrite | PDM13 | 0.684 | 142 | 44 |
| nitrite | PDM14 | 0.835 | 151 | 35 |
| nitrite | PDM21 | 0.036 | 151 | 35 |
| nitrite | PNT03 | 0.815 | 139 | 47 |
| nitrite | PXM03 | 0.875 | 153 | 33 |
| nitrate | ACM01 | 0.864 | 123 | 63 |
| nitrate | ACV02 | 0.871 | 157 | 29 |
| nitrate | AGB01 | 0.893 | 138 | 48 |
| nitrate | ENS09 | 0.856 | 137 | 49 |
| nitrate | PAR01 | 0.258 | 137 | 49 |
| nitrate | PAR19367 | 0.575 | 153 | 33 |
| nitrate | PDM12 | 0.958 | 150 | 36 |
| nitrate | PDM13 | 0.940 | 142 | 44 |
| nitrate | PDM14 | 0.948 | 151 | 35 |
| nitrate | PDM21 | -0.088 | 151 | 35 |
| nitrate | PNT03 | 0.941 | 139 | 47 |
| nitrate | PXM03 | 0.955 | 153 | 33 |
| acetate | AC | 0.685 | 282 | 294 |
| acetate | BA | 0.751 | 381 | 195 |
| acetate | BC | 0.885 | 445 | 131 |
| acetate | BF | 0.919 | 467 | 109 |
| acetate | BH | 0.863 | 396 | 180 |
| acetate | BL | 0.912 | 433 | 143 |
| acetate | BO | 0.849 | 437 | 139 |
| acetate | BP | 0.687 | 418 | 158 |
| acetate | BT | 0.911 | 469 | 107 |
| acetate | BU | 0.840 | 440 | 136 |
| acetate | BV | 0.828 | 394 | 182 |
| acetate | BY | 0.582 | 434 | 142 |
| acetate | CA | 0.847 | 417 | 159 |
| acetate | CC | 0.608 | 189 | 387 |
| acetate | CG | 0.853 | 423 | 153 |
| acetate | CH | 0.859 | 423 | 153 |
| acetate | DF | 0.858 | 384 | 192 |
| acetate | DL | 0.759 | 447 | 129 |
| acetate | DP | 0.830 | 419 | 157 |
| acetate | EL | 0.881 | 422 | 154 |
| acetate | ER | 0.651 | 193 | 383 |
| acetate | FP | 0.708 | 195 | 381 |
| acetate | PC | 0.869 | 398 | 178 |
| acetate | PJ | 0.832 | 403 | 173 |
| acetate | RI | 0.685 | 203 | 373 |
| starch degradation | B_cereus | 0.857 | 31 | 22 |
| starch degradation | B_megaterium | 0.865 | 25 | 28 |
| starch degradation | B_mojavensis | 0.833 | 25 | 28 |

Continued on next page

Supplementary Table 2: Leave-one-out performances

| dataset | training species | pearson r | n_train | n_test |
|---|---|---|---|---|
| starch degradation | B_subtilis | 0.837 | 31 | 22 |
| starch degradation | B_thuringiensis | 0.887 | 25 | 28 |
| starch degradation | P_polymyxa | 0.218 | 25 | 28 |
| butyrate | AC | 0.399 | 282 | 294 |
| butyrate | BA | 0.918 | 381 | 195 |
| butyrate | BC | 0.922 | 445 | 131 |
| butyrate | BF | 0.951 | 467 | 109 |
| butyrate | BH | 0.871 | 396 | 180 |
| butyrate | BL | 0.934 | 433 | 143 |
| butyrate | BO | 0.936 | 437 | 139 |
| butyrate | BP | 0.942 | 418 | 158 |
| butyrate | BT | 0.957 | 469 | 107 |
| butyrate | BU | 0.938 | 440 | 136 |
| butyrate | BV | 0.938 | 394 | 182 |
| butyrate | BY | 0.908 | 434 | 142 |
| butyrate | CA | 0.947 | 417 | 159 |
| butyrate | CC | 0.837 | 189 | 387 |
| butyrate | CG | 0.934 | 423 | 153 |
| butyrate | CH | 0.924 | 423 | 153 |
| butyrate | DF | 0.934 | 384 | 192 |
| butyrate | DL | 0.889 | 447 | 129 |
| butyrate | DP | 0.840 | 419 | 157 |
| butyrate | EL | 0.946 | 422 | 154 |
| butyrate | ER | 0.827 | 193 | 383 |
| butyrate | FP | 0.840 | 195 | 381 |
| butyrate | PC | 0.950 | 398 | 178 |
| butyrate | PJ | 0.939 | 403 | 173 |
| butyrate | RI | 0.814 | 203 | 373 |
| lactate | AC | 0.384 | 282 | 294 |
| lactate | BA | 0.869 | 381 | 195 |
| lactate | BC | 0.905 | 445 | 131 |
| lactate | BF | 0.910 | 467 | 109 |
| lactate | BH | 0.916 | 396 | 180 |
| lactate | BL | 0.932 | 433 | 143 |
| lactate | BO | 0.934 | 437 | 139 |
| lactate | BP | 0.877 | 418 | 158 |
| lactate | BT | 0.880 | 469 | 107 |
| lactate | BU | 0.917 | 440 | 136 |
| lactate | BV | 0.846 | 394 | 182 |
| lactate | BY | 0.884 | 434 | 142 |
| lactate | CA | 0.924 | 417 | 159 |
| lactate | CC | 0.725 | 189 | 387 |
| lactate | CG | 0.892 | 423 | 153 |
| lactate | CH | 0.900 | 423 | 153 |
| lactate | DF | 0.894 | 384 | 192 |
| lactate | DL | 0.824 | 447 | 129 |
| lactate | DP | 0.578 | 419 | 157 |
| lactate | EL | 0.883 | 422 | 154 |
| lactate | ER | 0.640 | 193 | 383 |
| lactate | FP | 0.727 | 195 | 381 |
| lactate | PC | 0.917 | 398 | 178 |
| lactate | PJ | 0.925 | 403 | 173 |

Continued on next page

Supplementary Table 2: Leave-one-out performances

| dataset | training species | pearson r | n_train | n_test |
|---|---|---|---|---|
| lactate | RI | 0.726 | 203 | 373 |
| *C. difficile* inhibition | BH | 0.871 | 97 | 83 |
| *C. difficile* inhibition | BO | 0.805 | 126 | 54 |
| *C. difficile* inhibition | BT | 0.749 | 134 | 46 |
| *C. difficile* inhibition | BU | 0.800 | 114 | 66 |
| *C. difficile* inhibition | BV | 0.820 | 131 | 49 |
| *C. difficile* inhibition | CA | 0.836 | 102 | 78 |
| *C. difficile* inhibition | CH | 0.780 | 93 | 87 |
| *C. difficile* inhibition | CS | 0.820 | 140 | 40 |
| *C. difficile* inhibition | DP | 0.798 | 69 | 111 |
| *C. difficile* inhibition | EL | 0.826 | 128 | 52 |
| *C. difficile* inhibition | ER | 0.845 | 124 | 56 |
| *C. difficile* inhibition | FP | 0.785 | 114 | 66 |
| *C. difficile* inhibition | PC | 0.820 | 109 | 71 |
| succinate | AC | 0.813 | 282 | 294 |
| succinate | BA | 0.824 | 381 | 195 |
| succinate | BC | 0.757 | 445 | 131 |
| succinate | BF | 0.635 | 467 | 109 |
| succinate | BH | 0.856 | 396 | 180 |
| succinate | BL | 0.782 | 433 | 143 |
| succinate | BO | 0.812 | 437 | 139 |
| succinate | BP | 0.881 | 418 | 158 |
| succinate | BT | 0.656 | 469 | 107 |
| succinate | BU | 0.694 | 440 | 136 |
| succinate | BV | 0.725 | 394 | 182 |
| succinate | BY | 0.777 | 434 | 142 |
| succinate | CA | 0.800 | 417 | 159 |
| succinate | CC | 0.750 | 189 | 387 |
| succinate | CG | 0.843 | 423 | 153 |
| succinate | CH | 0.802 | 423 | 153 |
| succinate | DF | 0.804 | 384 | 192 |
| succinate | DL | 0.817 | 447 | 129 |
| succinate | DP | 0.890 | 419 | 157 |
| succinate | EL | 0.746 | 422 | 154 |
| succinate | ER | 0.765 | 193 | 383 |
| succinate | FP | 0.786 | 195 | 381 |
| succinate | PC | 0.704 | 398 | 178 |
| succinate | PJ | 0.734 | 403 | 173 |
| succinate | RI | 0.792 | 203 | 373 |

| datasets | number of folds |
|---|---|
| acetate, butyrate, lactate, succinate (Clark et al. 2021) | 8 |
| nitrate and nitrite (Gowda et al 2022) | 6 |
| *C. difficile* inhibition (Hromada et al 2021) | 9 |
| starch degradation (Sanchez-Gorostiaga et al 2019) | 9 |

Supplementary Table 3: **Number of folds used in $k$-fold cross-validation for each dataset.**

| Co-A transferase (pwy 1) | butyrate kinase (pwy 2) | lactate to pyruvate (pwy 3) | acetate ($y_1$) | butyrate ($y_2$) | lactate ($y_3$) |
|---|---|---|---|---|---|
| 0 | 0 | NA | $U[-1,1]$ | $-1.3$ | $U[-1,1]$ |
| 0 | 1 | 0 | $U[-1,1]$ | $U[-0.5,0]$ | $U[-1,1]$ |
| 0 | 1 | 1 | $U[-1,1]$ | $U[-0.5,0]$ | $-y_2$ |
| 1 | NA | 1 | $-y_2$ | $U[0,2]$ | $-y_2$ |
| 1 | 1 | 0 | $-y_2$ | $U[0,0.5]$ | $U[-1,1]$ |
| 1 | 0 | 0 | $-y_2$ | $U[0,0.5]$ | $U[-1,1]$ |

Supplementary Table 4: **Acetate, butyrate, and lactate produced synthetic species monoculture with different combinations of driver genetic feature.** 'NA' indicates the pathway can be either present or absent; 0 (1) indicates the pathway is absent (present), respectively. Recall that all community functional data were standardized to have zero mean and unit variance. A uniform distribution on the interval $(x, y)$ is denoted by $U[x, y]$. When pwy 1 and pwy 2 are both absent $y_2$ is assigned to -1.3 to reflect the butyrate amount corresponding to 0 mM standardized by experimental mean and standard deviation.
